# CEP44 is required for maintaining centriole duplication and spindle integrity

**DOI:** 10.1101/2023.11.20.567852

**Authors:** Donghui Zhang, Wenlu Wei, Xiaopeng Zou, Hui Meng, Fangyuan Li, Minjun Yao, Junling Teng, Ning Huang, Jianguo Chen

## Abstract

In animal cells, the centrosome, consisting of two centrioles, duplicates only once per cell cycle for bipolar spindle formation. Defective centriole duplication results in abnormal spindle formation and chromosome missegregation, which is closely linked to tumor growth. However, the molecular mechanisms licensing only one centriole duplication cycle within a cell cycle are less well known. Here we found that CEP44 is negatively correlated with breast carcinoma. CEP44, jointly with CEP57 and CEP57L1, maintains centriole engagement in the interphase to ensure centriole duplication once per cell cycle. Depletion of CEP44 leads to centriole overduplication because of premature centriole disengagement, and multipolar spindle formation. Additionally, CEP44 is phosphorylated by Aurora A at the G2/M phase to facilitate spindle localization and maintain spindle integrity. Collectively, our results show the function of CEP44 in spindle formation by preventing centriole overduplication and maintaining spindle integrity, and CEP44 may serve as a potential marker for breast carcinoma prognosis.

## Introduction

The centrosome, at the center of which is a pair of centrioles, is surrounded by pericentriolar material (PCM) and centriolar satellites, and acts as the major microtubule organizing center (MTOC) in mammalian cells (1). In mitosis, the two centrosomes move to the opposite side in a cell and function in the nucleation of microtubules important for forming the bipolar spindle to ensure even segregation of genetic material. Therefore, centriole duplication, occurring only once per cell cycle, is strictly initiated through assembly of the procentriole at the proximal parts of preexisting centrioles during the S phase (2–4). Anomalies in centriole duplication caused by improper centrosome number control lead to disrupted spindle formation and chromosomal segregation errors, and eventually culminate in cancer and other developmental diseases (5–8).

The molecular mechanisms controlling only one centriole duplication cycle within a cell cycle are not well understood, despite significant advancements in understanding copy number control of centriole duplication in the S phase (3). Maintenance of centriole engagement in the interphase has been reported to be fundamental for ensuring that centriole duplication occurs only once per cell cycle (9). Immediately after centriole duplication, the newly formed daughter centriole is perpendicularly engaged with preexisting centrioles until late mitosis (centrosome engagement). After mitotic exit, the close-connected mother and daughter centrioles are separated (centrosome disengagement), and this separation allows for centriole duplication in the next cell cycle. When premature centriole disengagement happens as early as the S phase, centrioles are reduplicated, producing numerous centrioles within the same cell cycle (10).

It has been reported that PCM plays a pivotal role not only in microtubule nucleation, but also in modulation of centriole engagement and disengagement both in mitosis and interphase (11). Pericentrin (PCNT), a PCM scaffold protein, is critical for ensuring the maintenance of centriole engagement in mitosis. For normal centriole disengagement during mitotic exit, PCNT is specifically cleaved by separase to disassemble the expanded PCM (12). PLK1-dependent phosphorylation of PCNT is a prerequisite for the cleavage of PCNT by separase (13).

CEP57, a PCM protein, is closely associated with tumor development and variegated aneuploidy syndrome (14, 15). As a crucial binding partner of PCNT, CEP57 serves as a PCM organization modulator (16); CEP57 depletion causes premature centriole disengagement in mitosis (17). Notably, co-depletion of CEP57 with its paralog CEP57L results in premature centriole disengagement in interphase, resulting in centriole reduplication and disturbed chromosome segregation by amplified centrioles (9). Thus, centriole engagement in interphase is of much greater importance than that in mitosis for centriole duplication and proper chromosome segregation. However, the molecular mechanisms underlying centriole engagement in interphase are largely unknown, although there has been some progress in understanding the basis of centriole engagement in mitosis.

A centrosomal protein of 44 kDa (CEP44) was first identified as a centrosome protein, and probable spindle protein, via complementary proteomics (18). It has been reported that CEP44 is a centrosome linker protein that functions in centrosome cohesion (19). Recent studies also demonstrate that CEP44, as a centriole luminal protein, plays a vital role in the centriole-to-centrosome conversion program by promoting the formation of structurally-intact centrioles with a 9-fold symmetric triplet microtubule structure (20). In this study, we show that the expression of CEP44 is negatively correlated with breast carcinogenesis. CEP44 ensures centriole duplication once per cell cycle by maintaining centriole engagement with CEP57 and CEP57l1 in interphase. In mitosis, CEP44 is required for spindle integrity and is activated by Aurora A-mediated phosphorylation.

## Results

### CEP44 is negatively correlated with breast carcinogenesis

Several studies have conclusively demonstrated that certain centrosome proteins act as powerful tumor suppressors or activators (21–24). The centrosomes in breast cancer cells show characteristic structural and functional aberrations attributed to dysregulation of important centrosome or non-centrosome proteins (8, 25–31). To reveal the mechanisms underlying intracellular centrosome structure and function in breast cancer cells, we conducted a search for new centrosome proteins that might function in intrinsic mechanisms of centrosome structure and function in breast cancer cells. Intriguingly, MDA-MB-436 cells showed a nearly total loss of CEP44 protein expression and centrosome localization (Figure 1A-C). Human CEP44 consists of 390 amino acids (aa) containing two short coiled-coil domains in the middle region and C-terminus (Figure 1–figure supplement 1A). To confirm the specificity of the antibodies against CEP44, CEP44 was depleted by CEP44 siRNAs and detected by immunoblots. Both our homemade and commercial antibodies recognized a band at approximately 45 kDa by immunoblotting in HeLa cells, and this band was specifically abolished by siRNAs targeting CEP44 (Figure 1–figure supplement 1B and C).

**Figure 1.**
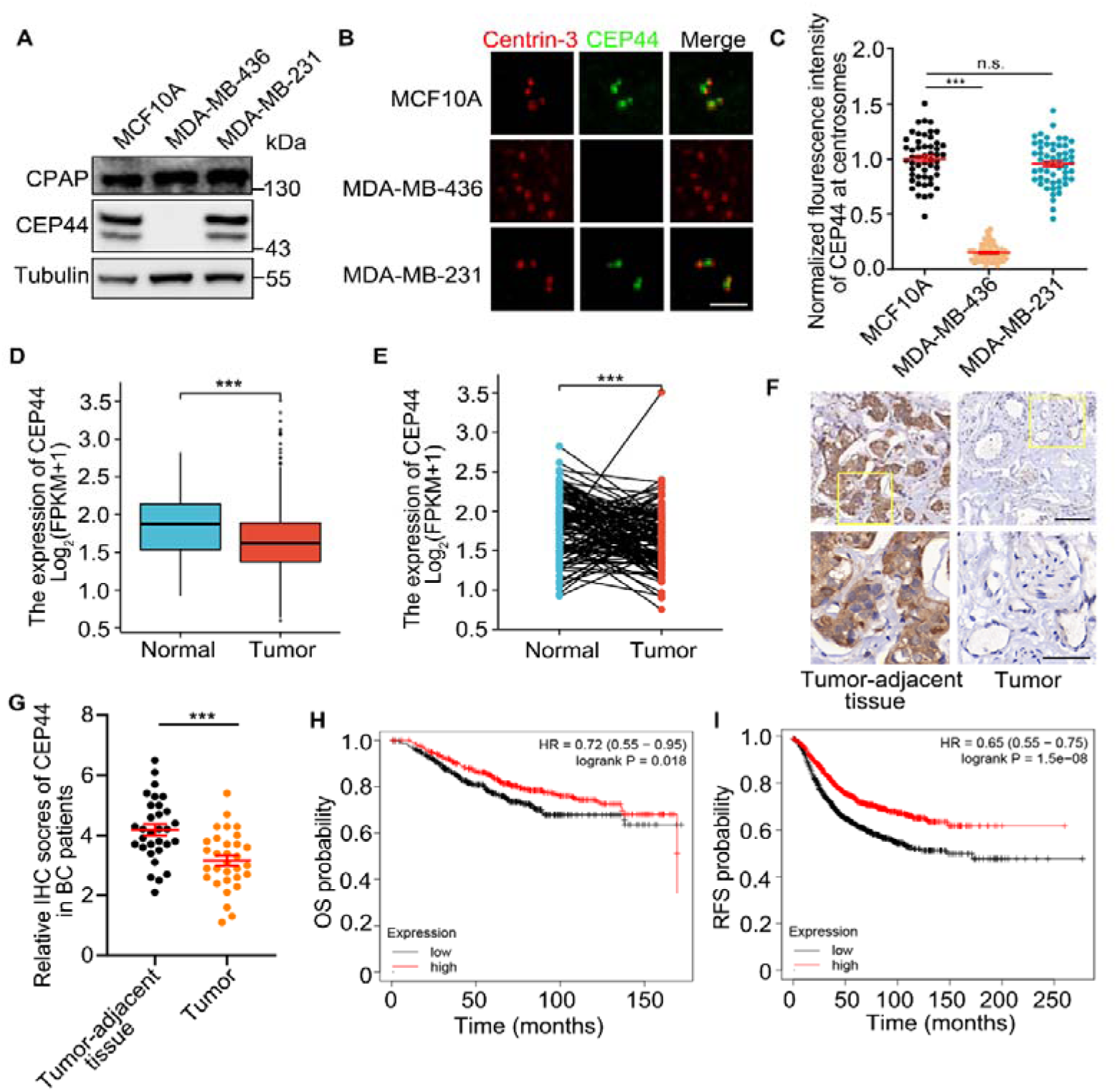
Downregulation of CEP44 in breast cancer correlates with poor prognosis. (A) Immunoblots of CPAP and CEP44 in MCF10A, MDA-MB-436 and MDA-MB-231 cells. Tubulin was used as a loading control. (B) Immunostaining of Centrin-3 (red) and CEP44 (green) in MCF10A, MDA-MB-436 and MDA-MB-231 cells. (C) Quantification of the fluorescence intensity of CEP44 in the cells from (B). (D) CEP44 transcript levels in normal human mammary tissues and breast carcinoma samples based on data sets from TCGA. (E) CEP44 transcript levels in paired tumor and tumor-adjacent tissue samples based on data sets from TCGA. (F-G) Representative images (F) of IHC staining and the relative IHC scores (G) of CEP44 in breast cancer tissues and adjacent normal tissues. (H-I) Overall survival (OS) (H) and recurrence free survival (RFS) (I) were compared between breast cancer patients with high and low CEP44 expression using Kaplan-Meier Plotter. For C, error bars represent the means ± S.E.M for three independent experiments. n.s., not significant, \*\*\**p* < 0.001, as determined using one-way ANOVA with Dunnett’s multiple comparisons test. For D, E and G, error bars represent the means ± S.E.M for three independent experiments. \*\*\**p* < 0.001, as determined using Student’s t-test. Bars: 2 μm (B), 100 μm (F, upper), 50 μm (F, lower). **Figure 1-soure data 1** Information of collected breast cancer patients.

Moreover, the expression data obtained from TCGA showed lower expression of CEP44 mRNA in breast cancer patients in comparison with normal individuals (Figure 1D), as well as in breast cancer tissue in comparison with normal tissue (Figure 1E). Consistently, immunohistology analysis of CEP44 expression levels using human breast tissue arrays that included paired tumor and tumor-adjacent tissues showed that CEP44 was expressed at a low level in breast carcinoma samples (Figure 1F-G). Analysis of data from the cancer genome atlas program (TCGA) cohort also revealed that lower CEP44 expression in breast cancer patients was associated with worse overall survival (OS) and recurrence free survival (RFS) outcomes (Figure H-I). In conclusion, these observations suggest a vital role for CEP44 in breast carcinogenesis.

Detailed localization of CEP44 by stimulated emission depletion (STED) microscopy showed that CEP44 formed a small ring-like structure, enclosed by the circle marked by polyglutamylated and acetylated tubulin, a centriole wall marker (from the top view), and localized tightly along the inner side of the circle (from the side view) (Figure 1–figure supplement 1D-G), suggesting that CEP44 is a luminal centriole protein as previously reported (20). 3D-structured illumination microscopy (3D-SIM) showed that CEP44 colocalized with C-Nap1, a marker of the centrosome linker at the proximal ends, more closely than it colocalized with CEP164 or GFP-Centrin-3 (both markers of the distal ends of centrioles), suggesting that CEP44 is localized at the proximal halves of centrioles. Co-staining of CEP44 with CEP152 (localized to the proximal halves of centrioles) and CEP192 (localized along the entire walls of centrioles) (32) revealed that CEP44 was partly colocalized with CEP192 and encircled by CEP152 (Figure 1–figure supplement 1H), demonstrating that CEP44 is localized to the proximal halves of centrioles.

### CEP44 is a negative regulator of centriole duplication

As reported previously (33, 34), centrioles are remarkably overduplicated in breast cancer cells such as MDA-MB-436, MDA-MB-231, SKBR3 and HCC1806 cells, but not in the spontaneously immortalized, non-malignant breast cells MCF10A (Figure 2A). In particular, centriole overduplication was found to be particularly severe in the MDA-MB-436 cells (approximately 80%) (Figure 1B and Figure 2B), suggesting that it CEP44 may play critical role in regulating centriole duplication in breast cancer cells.

**Figure 2.**
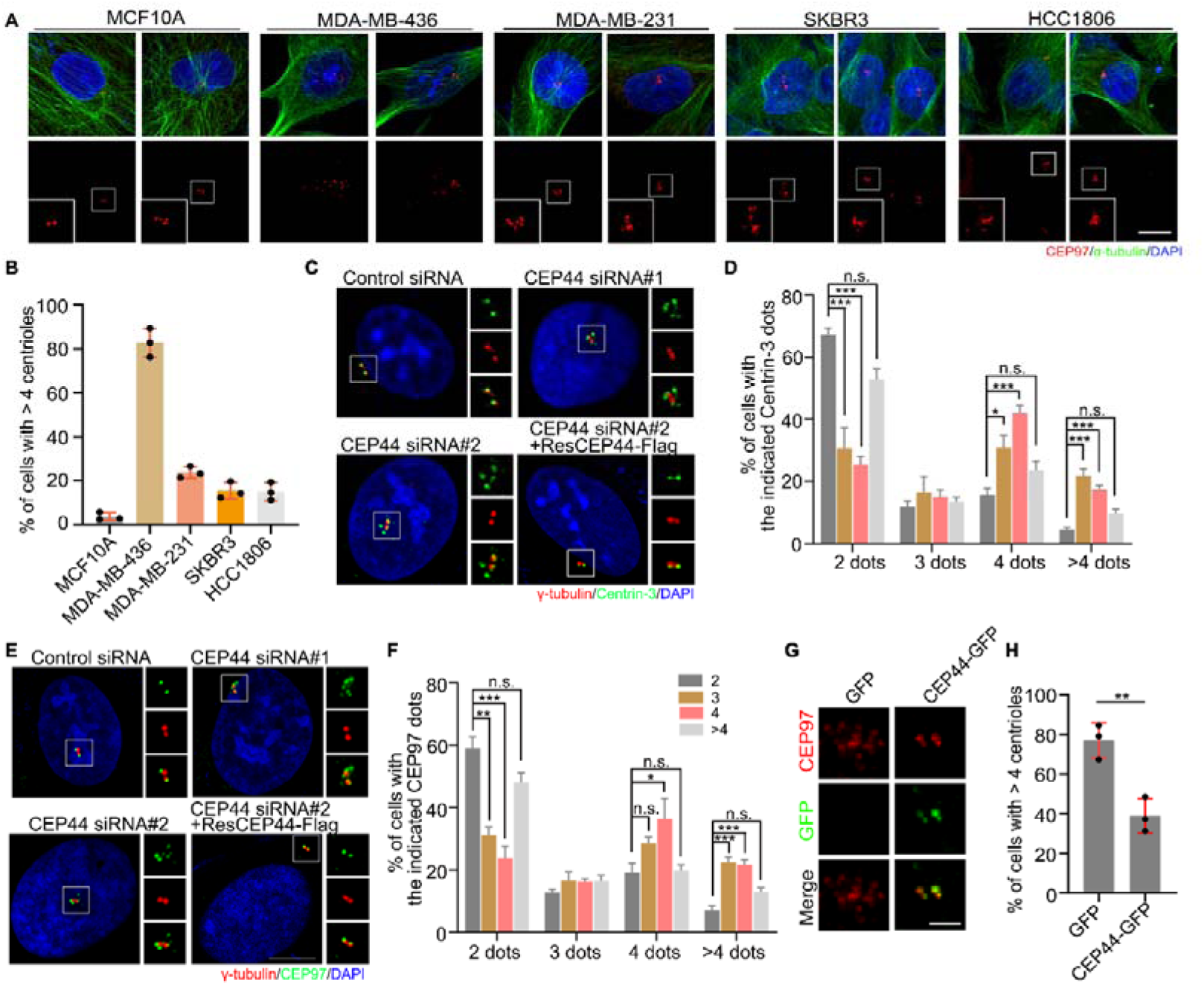
Depletion of CEP44 results in centriole overduplication. (A) Immunostaining of CEP97 (red) and α-tubulin (green) in MCF10A, MDA-MB-436, MDA-MB-231, SKBR3 and HCC1806 cells. DNA was stained with DAPI (blue). (B) Quantification of the percentage of cells with >4 centrioles from (A). (C-F) Immunostaining of Centrin-3 (green) (C) or CEP97 (green) (E) with γ-tubulin (red) in HeLa cells transfected with control-or CEP44-siRNA. DNA was stained with DAPI (blue). Quantification of the percentage of cells with the indicated number of Centrin-3 dots (D) and CEP97 dots (F) from (C and E). (G) Immunostaining of CEP97 (red) in CEP44-GFP-overexpressed MDA-MB-436 cells. (H) Quantification of the percentage of cells with more than 4 centrioles from (G). For B, D, F and H, error bars represent the means ± S.E.M for three independent experiments. n.s., not significant, \**p* < 0.05, \*\**p* < 0.01, \*\*\**p* < 0.001, as determined using one-way ANOVA with Dunnett’s multiple comparisons test (D-F) or Student’s t-test (H). Bars: 10 μm (A), 5 μm (C, E), 2 μm (G).

To investigate the function of CEP44 in centriole duplication, we performed siRNA knockdown experiments with two CEP44-targeting siRNAs in HeLa cells and observed remarkably increased centriole amplification (more than four centrioles marked by CEP97 or Centrin-3), which was rescued by overexpressing siRNA-resistant CEP44 (Figure 2C-F).

To investigate whether the increased number of centrioles in MDA-MB-436 cells is attributed a loss of CEP44, centriole numbers in MDA-MB-436 cells were recalculated after CEP44 ectopic overexpression. The fraction of cells showing centriole overduplication significantly decreased to approximately 40% in CEP44 overexpressed MDA-MB-436 cells (Figure 2G-H), demonstrating that CEP44 plays an important role in controlling centriole duplication in MDA-MB-436 cells.

### CEP44 maintains centriole engagement

To dissect the mechanism of centriole overduplication after CEP44 depletion, we observed centrosome phenotypes. Firstly, we generated CEP44 knockout HeLa cell lines using a CRISPR-Cas9 system (Figure 3–figure supplement 1A-B). CEP44-depleted cells showed increased distance (>0.75 μm) between newly formed daughter centriole and old centriole after centriole duplication in interphase and mitosis (Figure 3A-C), indicating that precocious centriole disengagement occurred after CEP44 depletion. To exclude the possibility that the four separated centrioles observed in interphase after CEP44 depletion resulted from failed cytokinesis, we determined the number of old mother centrioles; if cytokinesis failed, then two old mother centrioles were expected. Immunostaining with two markers of old mother centrioles, CEP164 and ODF2, revealed the presence of only one dot marked by CEP164 and ODF2 in ∼70% of CEP44-depleted cells with four separated or amplified centrioles (Figure 3–figure supplement 1C-D). These findings suggest that the separated centrioles were likely caused by premature centriole disengagement instead of cytokinesis failure.

**Figure 3.**
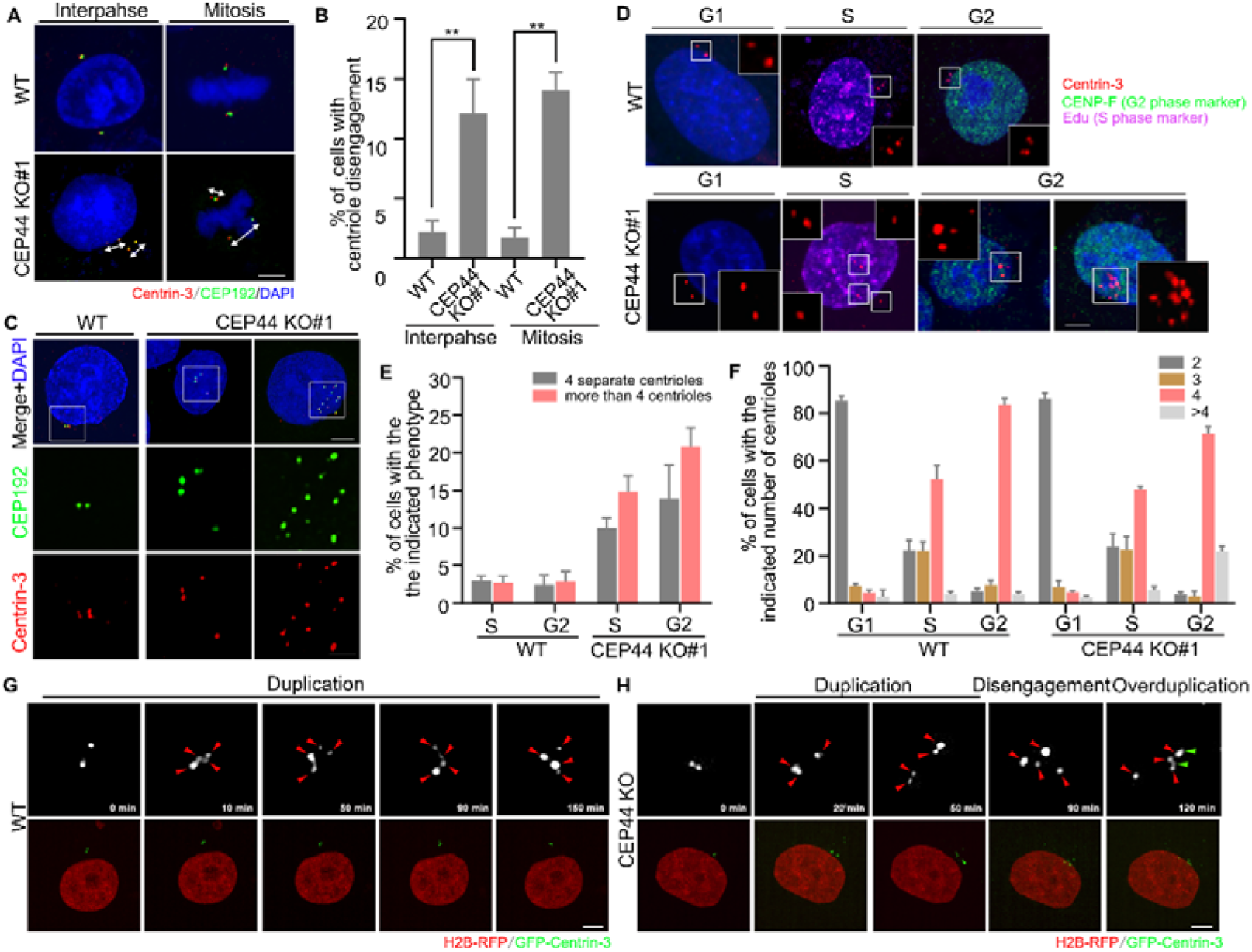
Depletion of CEP44 results in premature centriole disengagement. (A) Immunostaining of Centrin-3 (red) and CEP192 (green) in wild-type (WT) and CEP44 knockout (KO) HeLa cells. DNA was stained with DAPI (blue). (B) Quantification of the percentage of cells with centriole disengagement from (A). (C) Immunostaining of CEP192 (green) and Centrin-3 (red) in WT and CEP44 KO HeLa cells. DNA was stained with DAPI (blue). (D) Immunostaining of CENP-F (green) and Centrin-3 (red) in WT and CEP44 KO HeLa cells treated with EdU (S phase marker; purple) for 30 min before fixation. DNA was stained with DAPI (blue). (E-F) Quantification of the percentage of cells with the indicated phenotype (E) and number of centrioles (F) from (D). (G-H) Time-lapse images of WT (G) and CEP44 KO (H) HeLa cells stably expressing H2B-RFP and GFP-Centrin-3. Red arrowheads indicate centrioles after the first duplication. Green arrowheads indicate the overduplicated centrioles after the second overduplication. For B, E and F, error bars represent the means ± S.E.M obtained for three independent experiments. \*\**p* < 0.01, as determined using Student’s t-test. Bars: 5 μm (A, C upper, D, G and H), 2 μm (C, lower). **Figure 3–video 1** Video of WT HeLa cells stably expressing H2B-RFP and GFP-Centrin-3. **Figure 3–video 2** Video of CEP44 KO HeLa cells stably expressing H2B-RFP and GFP-Centrin-3.

Given that depletion of CEP44 resulted in both centriole disengagement and overduplication (Figure 2C-F, 3A-C and Figure 3–figure supplement 1E-H), we performed experiments to elucidate the mechanisms underlying these events. Firstly, we used an S phase marker (5-ethynyl-29-deoxyuridine, EdU) and a G2 phase marker (centromere protein F, CENP-F) to observe the occurrence of centriole disengagement and overduplication during particular stages of the cell cycle. In the G1 phase (Edu and CENP-F negative), CEP44-depleted cells and control cells both exhibited two centrioles (Figure 3D-F). However, CEP44-depleted cells exhibited a remarkably increased percentage of cells with four or more separated centrioles in the S phase (Edu positive and CENP-F negative) and G2 phase (Edu negative and CENP-F positive) (Figure 3D-F). These results verify that CEP44 depletion results in centriole disengagement as early as the S phase, and thus CEP44 knockout cells exhibit four or more separate centrioles. Notably, centriole overduplication also occurred as early as the S phase after CEP44 depletion. However, these results did not show the sequential order of centriole disengagement and overduplication caused by CEP44 depletion, as both of these events occurred in the S phase. Therefore, to address the link between these events, centriole dynamics were monitored in cells transfected with GFP-centrin-3 and H2B-RFP by tracing centrioles with time-lapse microscopy. In control cells, maintenance of centriole engagement was observed after centriole duplication, whereas CEP44 knockout cells showed premature centriole disengagement followed by centriole overduplication per cell cycle (Figure 3G-H), suggesting that premature centriole disengagement in interphase contributes to centriole overduplication. It is worth noting that centriole overduplication after premature centriole disengagement resulting from CEP44 depletion generally led to the formation of only one newly formed daughter centriole, instead of excessive centrioles, from each pre-existing centriole (Figure 3H).

As the daughter centriole is formed, PCM is recruited to it to ensure initiation of the first stage of centrosome duplication, referred to as centriole-to-centrosome conversion (CCC). It has been reported that CEP44 is required for CCC; therefore, it is conceivable that the CCC of precociously disengaged daughter centrioles was impaired in CEP44-depleted cells in interphase, which may have impaired centriole duplication, rather than inducing centriole overduplication, as shown by CEP44-depleted cells in our assay. To address this paradox, we co-stained γ-tubulin and centrin-3 in CEP44 knockout cells. CEP44-depleted cells with more than 2 γ-tubulin-positive centrioles were quite rare before the G2 phase, but they gradually became more abundant during the G2 phase, during which cells exhibited two dots with strong γ-tubulin staining, and one or two dots with weak γ-tubulin staining. The light dots stained by γ-tubulin most likely represent the disengaged daughter centrioles. After centriole duplication in CEP44-depleted cells, we also detected recruitment of γ-tubulin in the reduplicated centrioles (Figure 3–figure supplement 1I-J). These findings demonstrate that PCM is initially absent in the precociously disengaged daughter centrioles, and PCM components are gradually recruited to their surroundings, primarily during the G2 phase.

### CEP44 maintains centriole engagement via CEP57 and CEP57L1

To determine the molecular mechanism underlying centriole engagement regulation by CEP44, we searched for proteins capable of interacting with CEP44. Our immunoprecipitation assay showed that there was no interaction between CEP44 and several centriole engagement modulators, including PCNT, Sgo1 and SCC1, but the interaction was detected between CEP44 and CEP57/CEP57L (Figure 4A-B). Further immunoprecipitation assays showed that there was no interaction between CEP44 and several centriole luminal proteins known to be important for centriole duplication, including SAS6, STIL, PLK4 and CEP135. Given that CEP57/CEP57L1 is known to be involved in controlling centriole disengagement in interphase (9), these results suggest that the molecular mechanism by which CEP44 regulates centriole engagement is likely mediated by its interaction with CEP57/CEP57L1. Thus, we further examined the relationship among CEP44, CEP57, and CEP57L1. To determine whether CEP44 affects CEP57/CEP57L1 expression or localization, we first knocked down CEP44 expression in HeLa cells and observed CEP57/CEP57L1 expression and localization. Although depletion of CEP44 by two different siRNAs did not affect CEP57/CEP57L1 expression (Figure 4C), localization of CEP57/CEP57L1 at centrosomes was significantly reduced (Figure 4D-F). As previously reported (20), we also observed impaired γ-tubulin localization at centrosomes in CEP44-depleted cells (Figure 4D and Figure 4–figure supplement 1A-B). Conversely, depletion of CEP57 or CEP57L1 did not significantly affect CEP44 localization at centrosomes (Figure 4G-H and Figure 4–figure supplement 1A). Similar to the expression pattern of CEP44 in breast cancer patients vs normal individuals, the expression data obtained from TCGA showed lower expression of CEP57 mRNA in patients with breast cancer compared with normal individuals (Figure 4–figure supplement 1C), as well as in breast cancer tissue compared with normal tissue (Figure 4–figure supplement 1D). Taken together, these results suggest that CEP44 maintains centriole engagement by regulating CEP57/CEP57L1 localization at the centriole.

**Figure 4.**
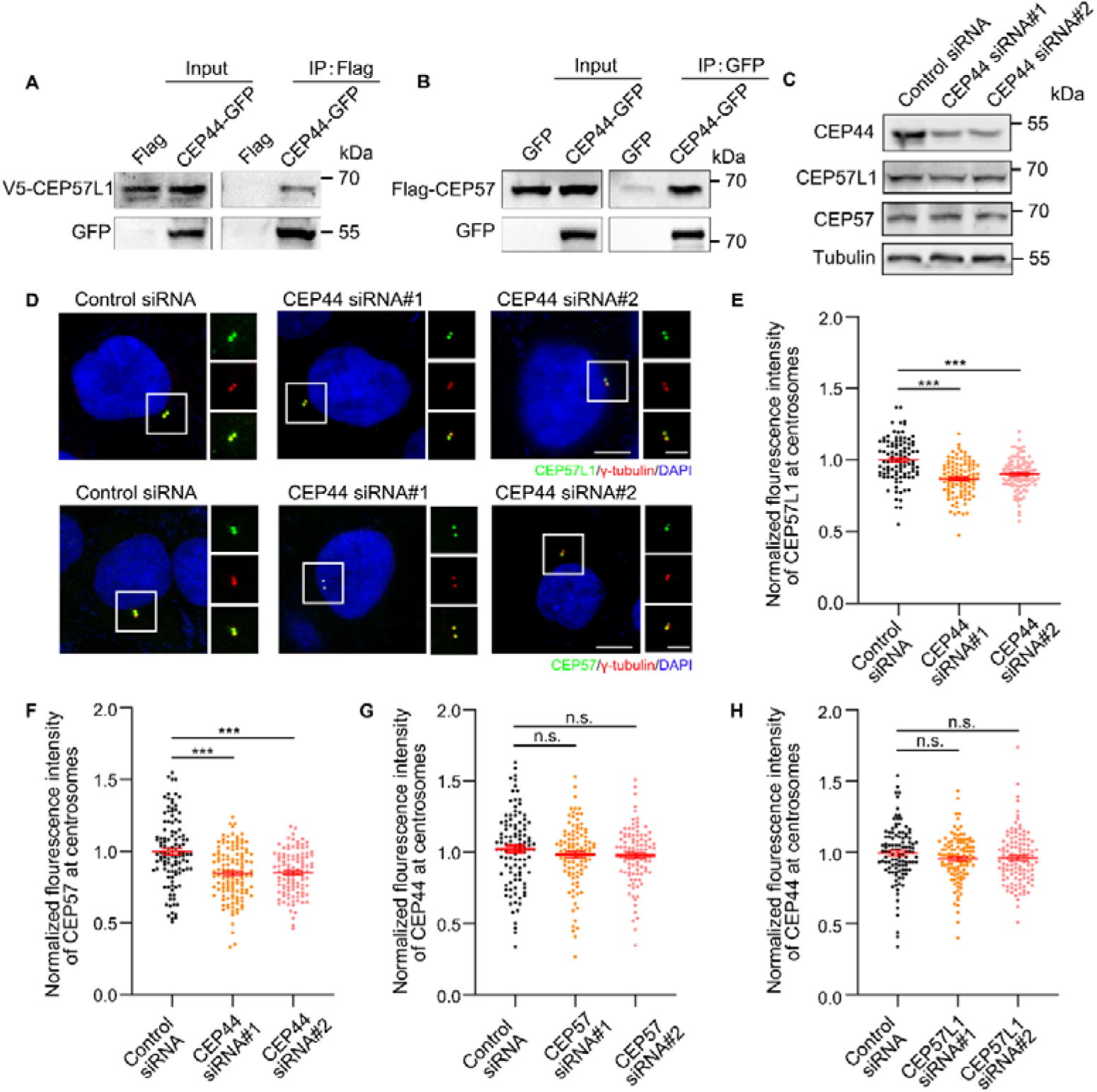
CEP44 recruits CEP57/CEP57L1 to the centriole. (A) Lysates of HEK293T cells co-overexpressing V5-CEP57L1 and Flag or CEP44-Flag were subjected to immunoprecipitation (IP) and immunoblotting with anti-Flag and anti-V5 antibodies. (B) Lysates of HEK293T cells co-overexpressing V5-CEP57 and Flag or CEP44-Flag were subjected to immunoprecipitation (IP) and immunoblotting with anti-Flag and anti-V5 antibodies. (C) Immunoblots of CEP44, CEP57L1 and CEP57 in control and CEP44-depleted HeLa cells. Tubulin was used as a loading control. (D) Immunostaining of CEP57L1 (green) or CEP57 (green) and γ-tubulin (red) in control and CEP44-depleted HeLa cells. DNA was stained with DAPI (blue). (E-F) Quantification of the fluorescence intensity of CEP57L1 (E) and CEP57 (F) in HeLa cells from (D). (G-H) Quantification of the fluorescence intensity of CEP44 in CEP57-depleted (G) and CEP57L1-depleted (H) HeLa cells. For E, F, G and H, error bars represent the means ± S.E.M for three independent experiments. n.s., not significant, \*\*\**p* < 0.001, as determined using one-way ANOVA with Dunnett’s multiple comparisons test. Bars: 5 μm (D).

### CEP44 depletion induced centriole overduplication leads to multipolar spindle formation

Next, we found that the percentage of multipolar and disorganized spindles increased remarkably in mitotic CEP44 knockout cells. This phenotype was nearly fully rescued by re-introduction of CEP44-Flag (Figure 5A-C). Similar phenotypes of increased percentages of multipolar and disorganized spindles in CEP44 knockdown HeLa cells were rescued by overexpressing siRNA-resistant CEP44 (Figure 5–figure supplement 1A-D).

**Figure 5.**
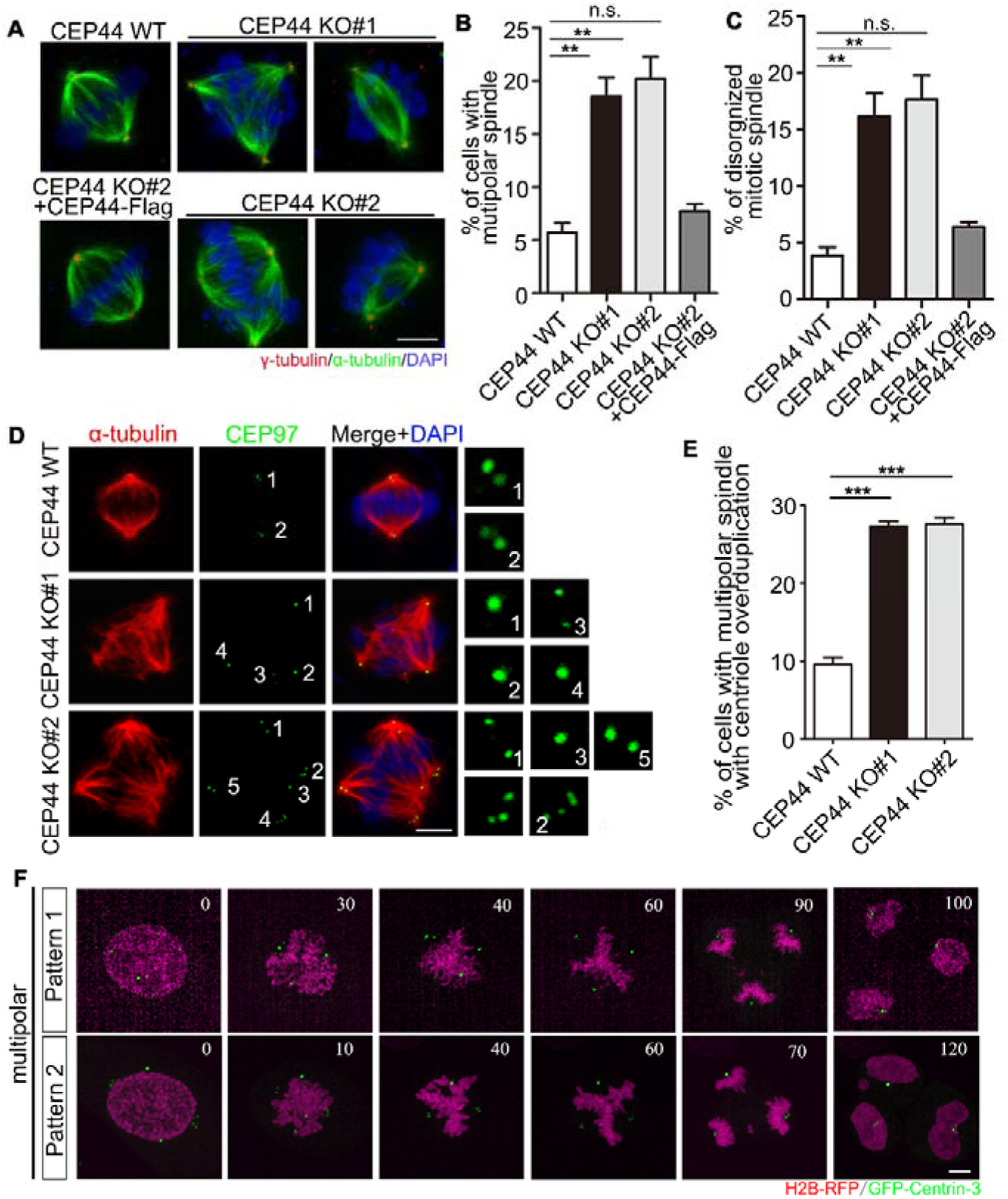
Depletion of CEP44 results in multipolar spindle formation. (A) Immunostaining of α-tubulin (green) and γ-tubulin (red) in wild-type (WT) or CEP44 knockout (KO) HeLa cells. DNA was stained with DAPI (blue). (B-C) Quantification of the percentage of cells with a multipolar spindle (B) or a disorganized spindle (C) from (A). (D) Immunostaining of CEP97 (green) and α-tubulin (red) in WT or CEP44 KO HeLa cells. DNA was stained with DAPI (blue). (E) Quantification of the percentage of cells with a multipolar spindle with centriole overduplication from (D). (F) Time-lapse images of mitotic CEP44 KO HeLa cells stably expressing H2B-RFP and GFP-Centrin-3. For B, C and E, error bars represent the means ± S.E.M obtained for three independent experiments. n.s., not significant, \*\**p* < 0.01, \*\*\**p* < 0.001, as determined using one-way ANOVA with Dunnett’s multiple comparisons test. Bars: 5 μm (A, D and F). **Figure 5–video 1** Video of multipolar spindle formation in mitotic CEP44 KO HeLa cells (pattern 1) stably expressing H2B-RFP and GFP-Centrin-3. **Figure 5–video 2** Video of multipolar spindle formation in mitotic CEP44 KO HeLa cells (pattern 2) stably expressing H2B-RFP and GFP-Centrin-3.

Given that depletion of CEP44 led to centriole amplification, we speculated that multipolar spindles in mitotic cells that formed after depletion of CEP44 were attributed to the presence of an elevated number of centrioles. To test our hypothesis, we stained mitotic HeLa cells with centriole marker CEP97 and calculated the centriole number. We observed an increased number of centrioles in multipolar spindles following depletion of CEP44 (Figure 5D-E and Figure 5–figure supplement 1E-F). These results indicate that depletion of CEP44 induced centrosome overduplication and abnormal spindle formation.

As mentioned above, CEP44-depleted HeLa cells exhibited a remarkably increased percentage of multipolar spindles (Figure 5A-B and Figure 5–figure supplement 1B-C). To determine whether centriole disengagement or/and centriole overduplication in interphase led to the increased percentage of multipolar spindles in CEP44 knockout cells, we transfected cells with GFP-centrin-3 and H2B-RFP and observed centrioles and chromosomes by live cell imaging. Based on the pattern of centriole disengagement and overduplication, we grouped cells into three categories in interphase: normal, premature centriole disengagement in interphase without centriole overduplication (pattern 1), or with centriole overduplication (pattern 2). In control cells, the two paired centrioles were engaged during the entire interphase (normal); while in CEP44 knockout cells, paired centrioles disengaged in interphase without centriole overduplication (pattern 1) or with centriole overduplication (pattern 2). Quantification of the percentage of multipolar spindles in cells showing pattern 1 or pattern 2 revealed that CEP44 knockout cells associated with centriole disengagement and centriole overduplication (pattern 2) exhibited the highest frequency of multipolar spindles (Figure 5F and Figure 5–figure supplement 1G). These results suggest that the formation of multipolar spindles after CEP44 depletion is mainly attributed to centriole overduplication resulting from premature centriole disengagement.

### Aurora A phosphorylates CEP44 in mitosis

Apart from its localization at centrosomes, CEP44 was also identified as a probable spindle protein in a complementary proteomics study (18). Thus, we determined whether CEP44 localizes at spindles, and found that it was readily detected at mitotic spindles throughout mitosis (Figure 6A and Figure 6–figure supplement 1A). To determine the pattern of CEP44 protein level changes during the cell cycle, we synchronized HeLa cells with nocodazole treatment. CEP44 protein levels peaked in the enriched M phase population (Figure 6B), similar to the well-known mitotic regulator cyclin B, indicating that CEP44 plays a critical role in mitosis. Multiple kinases, including Aurora A, PLK1, NEK2A, and CDK1, are localized to the spindle or centrosome and serve to regulate spindle dynamics during mitosis (35–38). To test whether these kinases can phosphorylate CEP44 to exert their function in mitosis, we firstly co-overexpressed CEP44 with these mitosis-related kinases in HEK293T cells. The band representing CEP44 was shifted slightly when CEP44 was co-overexpressed with wild-type Aurora A, but not kinase-dead Aurora A (Figure 6C and Figure 6–figure supplement 1B). However, this shifted band was completely abolished when the cell lysate was pretreated with lambda protein phosphatase (λ-PPase) (Figure 6C). Moreover, treatment with Aurora A inhibitor MLN8237 also abolished the shifted CEP44 band at the G2/M phase (Figure 6D and Figure 6–figure supplement 1C), indicating that CEP44 is phosphorylated by Aurora A at the G2/M phase. Mass spectrometry analysis revealed three sites that might be phosphorylated by Aurora A: Ser192, Ser324 and Ser342. To confirm the CEP44 phosphorylation site for Aurora A, we constructed unphosphorylatable mutants in which Ser192, Ser324 or Ser342 was replaced by alanine (S192A, S324A or S342A) using site-directed mutagenesis. The shifted band of CEP44-S324A was barely detected in cells co-overexpressing CEP44 and Aurora A (Figure 6E), indicating that CEP44 Ser324 is the site of Aurora-A-mediated phosphorylation. Additionally, Aurora A is known to recognize the consensus sequence R-X-pS/T-L/V for phosphorylation (39–42), and Ser324 of CEP44 accords with the consensus sequence (Figure 6F), confirming that Ser324 of CEP44 is phosphorylated by Aurora A. We then raised a phospho-CEP44 antibody against CEP44 Ser324 and verified the specificity of the antibody through peptide competition assays (Figure 6–figure supplement 1D). A band was detected when nocodazole-treated HeLa cell protein samples were incubated with the phospho-CEP44 antibody, but no band was detected from the control HeLa cells (Figure 6G). Moreover, depletion of Aurora A resulted in the loss of the band detected with the phospho-CEP44 antibody in nocodazole-synchronized HeLa cells (Figure 6H). Collectively, these results suggest that Aurora A phosphorylates CEP44 at Ser324 in the G2/M phase.

**Figure 6.**
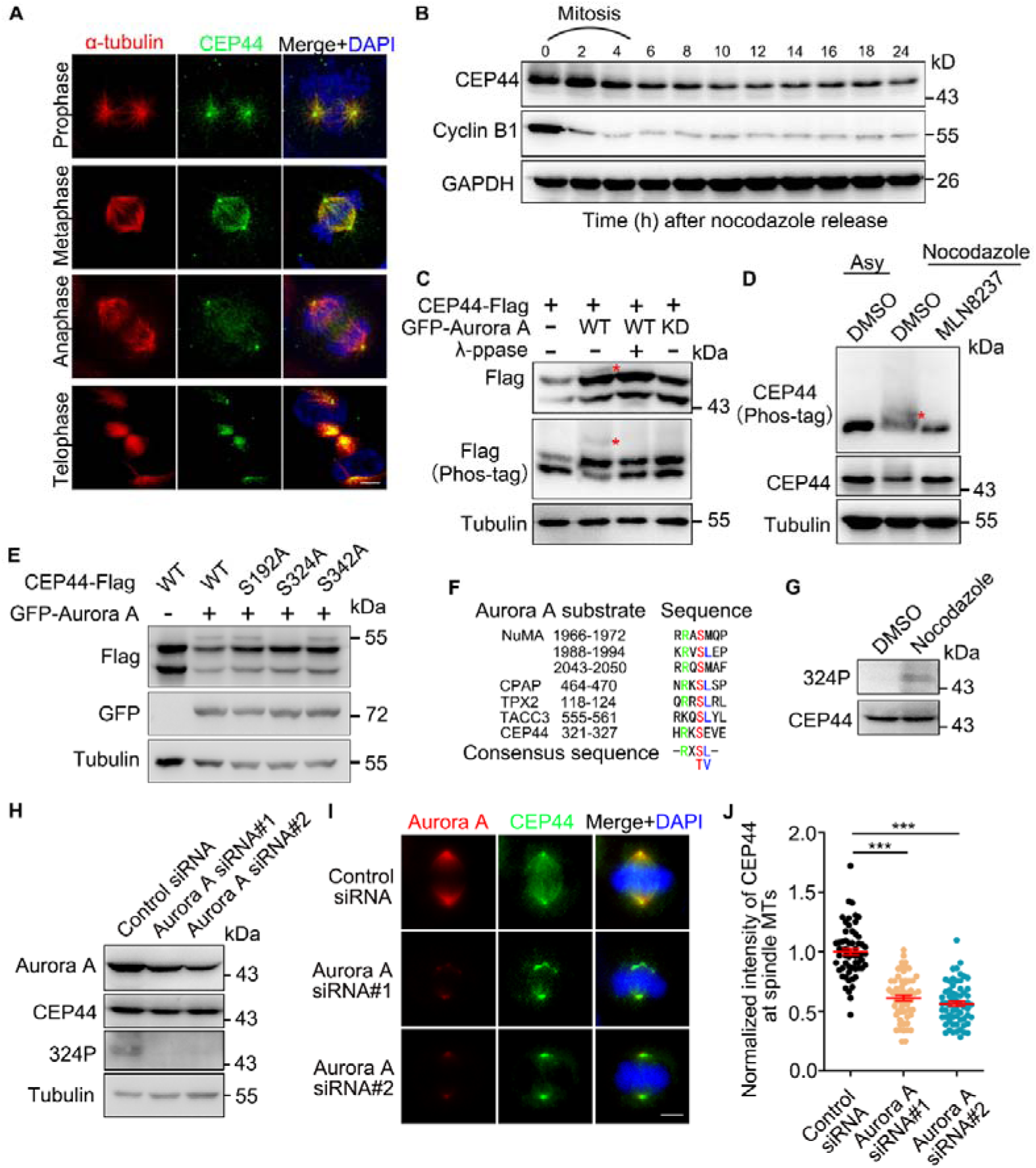
Aurora A phosphorylates CEP44 at S324 in the G2/M phase. (A) Immunostaining of α-tubulin (red) and CEP44 (green) in HeLa cells. DNA was stained with DAPI (blue). (B) Immunoblots of CEP44 and Cyclin B1 at the indicated time points after nocodazole release in HeLa cells. GAPDH was used as a loading control. (C) Immunoblots of lysates from HEK293T cells co-overexpressing the indicated proteins with or without λ-ppase treatment. Tubulin was used as a loading control. The red asterisks indicate the shifted band of CEP44. (D) Immunoblots of lysates from HEK293T cells with or without nocodazole and MLN8237 treatment. Tubulin was used as a loading control. The red asterisk indicates the shifted band of CEP44. (E) Immunoblots of lysates from HEK293T cells co-overexpressing the indicated proteins. Tubulin was used as a loading control. (F) The consensus sequence (R-X-pS/T-L/V) of the Aurora A substrate including NuMA, CPAP, TPX2, TACC3 and CEP44. (G) Immunoblots of lysates from HeLa cells with or without nocodazole treatment. (H) Immunoblots of the indicated proteins in control and Aurora A-depleted HeLa cells after nocodazole treatment. (I) Immunostaining of Aurora A (red) and CEP44 (green) in control and Aurora A-depleted mitotic HeLa cells. DNA was stained with DAPI (blue). (J) Quantification of the fluorescence intensity of CEP44 at spindle microtubules in mitotic cells from (I). For J, error bars represent the mean ± S.E.M. for three independent experiments. \*\*\**p* < 0.001, as determined using one-way ANOVA with Dunnett’s multiple comparisons test. Bars: 5 μm (A and I).

The results showing that CEP44 is highly expressed and phosphorylated during mitosis (Figure 6B-D) suggest that phosphorylation of CEP44 by Aurora A might alter its protein level. However, the CEP44 protein level did not change significantly after Aurora A knockdown in the G2/M phase (Figure 6H and Figure 6–figure supplement 1E). Next, we further examined the effect of Aurora-A-mediated phosphorylation on CEP44 localization. Immunostaining microscopy showed that mitotic Aurora A-depleted cells exhibited remarkably increased fluorescence intensity of CEP44 at the mitotic spindle pole, but less fluorescence intensity at the spindle, compared to the control cells (Figure 6I-J). This abnormal distribution of CEP44 in mitosis after Aurora A depletion was also detected in HeLa cells after treatment with MLN8237 (Figure 6–figure supplement 1F-H), suggesting that Aurora A-mediated phosphorylation of CEP44 regulates its localization between the spindle pole and spindle.

### CEP44 phosphorylation by Aurora A maintains spindle integrity

To clarify the physiological function of CEP44 and its phosphorylation at the spindle, we immunostained mitotic HeLa cells with α-tubulin to label spindle microtubules. Depletion of CEP44 significantly decreased the intensity of spindle microtubule staining, which was rescued after re-introduction of CEP44 (Figure 7A-B and Figure 7–figure supplement 1A-B), indicating that CEP44 regulates spindle microtubule stability. This finding suggests that CEP44 phosphorylation may be responsible for spindle microtubule stability, because CEP44 is phosphorylated in the G2/M phase. To determine whether CEP44 phosphorylation affects microtubule stability, CEP44 phosphorylation was mimicked in interphase HeLa cells by overexpressing phosphorylatable (CEP44-Flag S324D and S324E) or unphosphorylatable (CEP44-Flag S324A) mutants. Microtubule stability was not dramatically altered by overexpression of the phosphorylatable or unphosphorylatable mutants. However, after treatment with nocodazole for 5 min, HeLa cells overexpressing CEP44-Flag S324A exhibited more un-stabilized microtubules in comparison with HeLa cells overexpressing CEP44-Flag wild-type, S324D or S324E (Figure 7–figure supplement 1C), suggesting that CEP44 phosphorylation plays an important role in controlling microtubule stability. Similarly, in mitotic HeLa cells, ectopic overexpression of CEP44-Flag S324A remarkably impaired spindle microtubule stability (Figure 7C-E). Collectively, these results show the critical role of CEP44 phosphorylation in maintenance of microtubule stability and spindle structure. Notably, we found that CEP44-depleted cells showed weakened localization of γ-tubulin at the spindle (Figure 7F-G), which negatively affected spindle structure stability.

**Figure 7.**
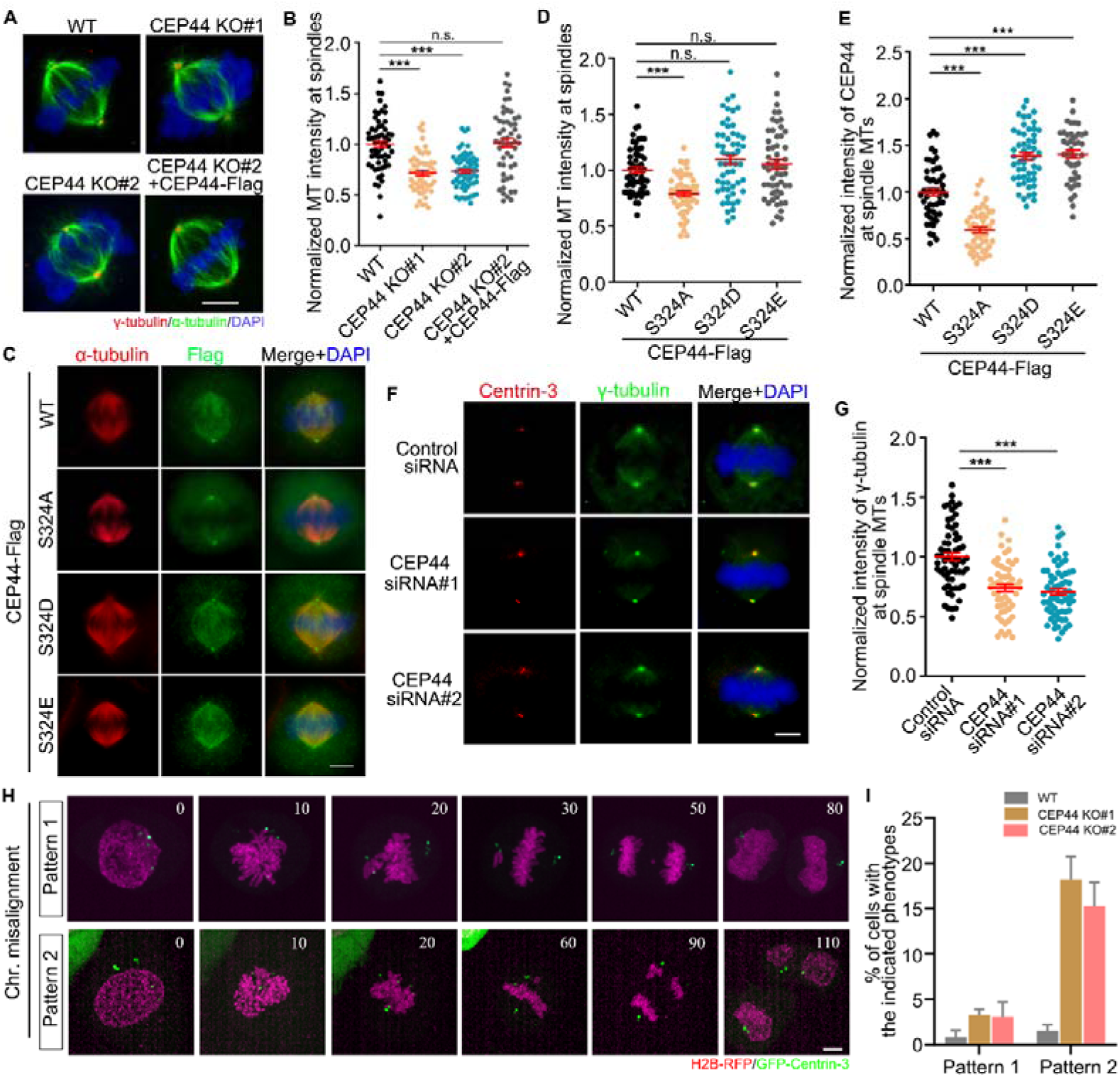
Phosphorylation of CEP44 by Aurora A maintains spindle integrity. (A) Immunostaining of γ-tubulin (red) and α-tubulin (green) in wild-type (WT) and CEP44 knockout (KO) HeLa cells. DNA was stained with DAPI (blue). (B) Quantification of fluorescence microtubule intensity in mitotic cells from (A). (C) Immunostaining of α-tubulin (red) and Flag (green) in mitotic HeLa cells expressing the indicated proteins. DNA was stained with DAPI (blue). (D-E) Quantification of the microtubule intensity at spindles (D) and intensity of CEP44 at spindle microtubules (E) in mitotic cells from (C). (F) Immunostaining of Centrin-3 (red) and γ-tubulin (green) in control and CEP44 knockdown HeLa cells. DNA was stained with DAPI (blue). (G) Quantification of the normalized intensity of γ-tubulin at spindle MTs in control and CEP44 knockdown HeLa cells. (H) Time-lapse images of mitotic CEP44 KO HeLa cells stably expressing H2B-RFP and GFP-Centrin-3. Chr, chromosome. (I) Quantification of the percentage of cells with the indicated phenotypes from (H). For B, D, E, G and I, error bars represent the means ± S.E.M for three independent experiments. n.s., not significant, \*\*\**p* < 0.001, as determined using one-way ANOVA with Dunnett’s multiple comparisons test. Bars: 5 μm (A, C, F and H). **Figure 7–video 1** Video of chromosome misalignment in mitotic CEP44 KO HeLa cells (pattern 1) stably expressing H2B-RFP and GFP-Centrin-3. **Figure 7–video 2** Video of chromosome misalignment in mitotic CEP44 KO HeLa cells (pattern 2) stably expressing H2B-RFP and GFP-Centrin-3.

Next, we investigated whether chromosome segregation errors occurred in mitotic CEP44-depleted cells due to impaired spindle structure (Figure 7A-B and 7F-G). CEP44-depleted cells showed a dramatically increased proportion of chromosome segregation errors, including misalignment and lagging chromosomes (Figure 7H-I), although bipolar spindle formation was accomplished eventually in some CEP44-depleted cells (Figure 7–figure supplement 1D). Therefore, CEP44 phosphorylation by Aurora A ensures normal spindle structure formation and chromosome segregation.

## Discussion

Our study indicates that centriole luminal protein CEP44 is a master regulator of centriole disengagement in interphase via its interaction with CEP57/CEP57L1, and its depletion results in premature centriole disengagement, centriole overduplication and multipolar spindle formation. Additionally, CEP44 is a spindle-localized protein that ensures spindle integrity, and this localization is modulated by Aurora A phosphorylation. Collectively, our results show that CEP44 maintains correct spindle formation through its localization at the centrosome and spindle (Figure 8).

**Figure 8.**
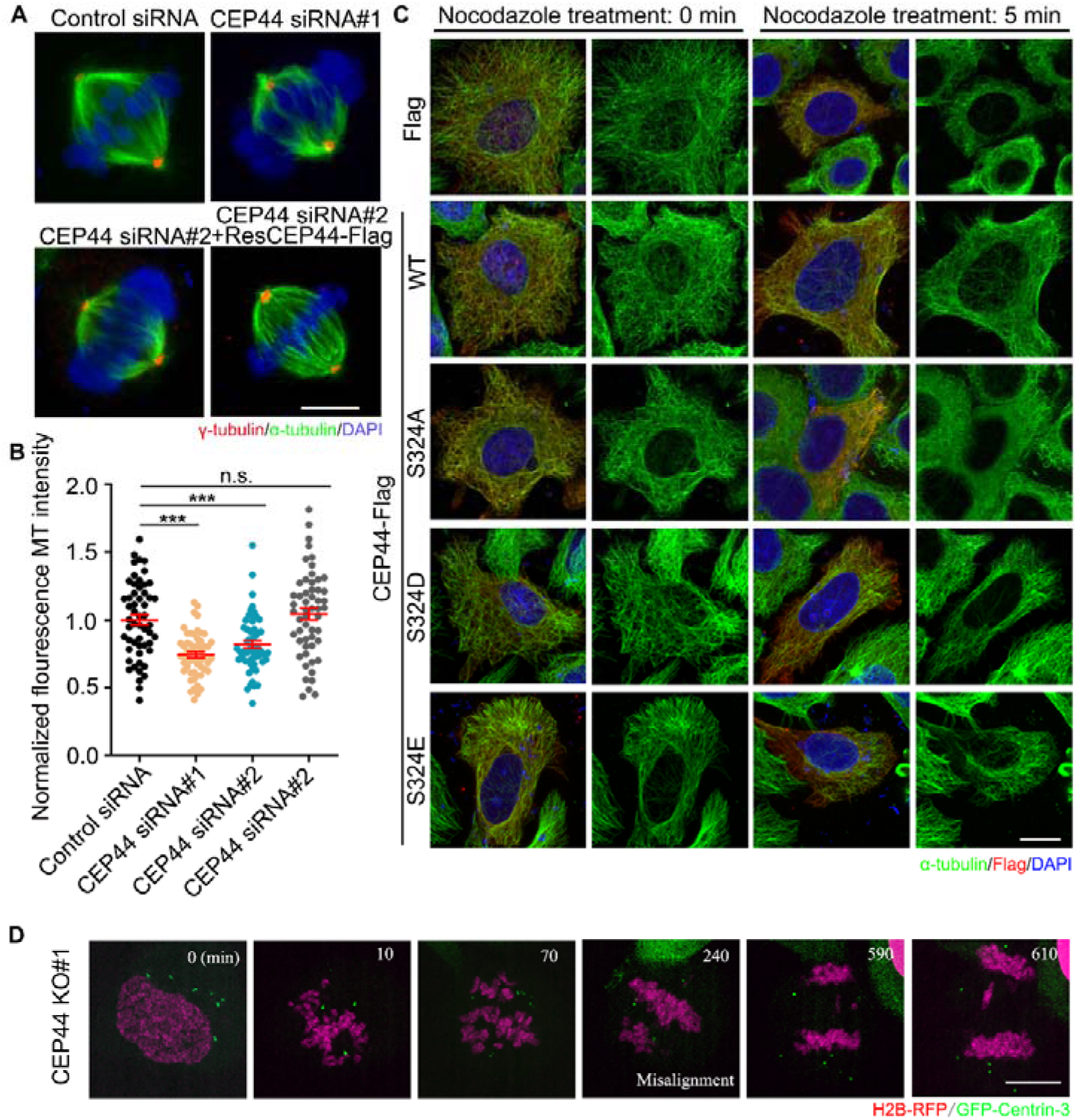
Schematic of the role of CEP44 in the cell cycle. In wild-type HeLa cells, CEP44 is localized at the proximal halves of the centriole lumen. For centriole duplication during the S phase, CEP44 exists in the lumen of the procentrioles to maintain disengagement between the old centrioles and the procentrioles. During mitosis, CEP44 translocates to the spindle due to Aurora A phosphorylation at serine 324 to ensure spindle integrity. After depletion of CEP44, premature centriole disengagement occurs following centriole duplication in the S phase, which licenses centriole overduplication as early as the S phase. The supernumerary centrioles result in multipolar spindle formation during mitosis. In addition, a lack of CEP44 localization at the spindle impairs spindle integrity, leading to an increase in the percentage of disorganized spindles. Notably, after Aurora A depletion or inhibition, unphosphorylated CEP44 fails to translocate to spindle microtubules and accumulates at the spindle pole, leading to impaired spindle integrity.

In contrast to the effects of co-depletion of PCM proteins CEP57 and CEP57L1, depletion of CEP44 alone was sufficient to result in premature centriole disengagement in interphase. A previous study reported that depletion of CEP44 impaired centrosome localization of PCM proteins, such as CEP152 and CEP192, which are required for centriole duplication (20). However, we showed centriole overduplication caused by premature centriole disengagement in CEP44-depleted cells, indicating that the effect of premature centriole disengagement in interphase, which accelerates centriole overduplication, overrides the effect of defective PCM recruitment (impaired CCC), which impedes centriole duplication, in the context of CEP44 depletion. These findings suggest that the residual PCM around prematurely disengaged centrioles is sufficient to induce centriole duplication after CEP44 deficiency. Collectively, our findings confirm that centriole disengagement is a critical prerequisite for a subsequent round of centriole duplication, indicating the fundamental role of centriole engagement/disengagement in modulating centriole duplication.

The centriole number is controlled based on two distinct rules to coordinate multiple cellular processes (2, 4); the first rule ensures that centrosomes duplicate once and only once per cell cycle (cell cycle control), and the second rule stipulates that only one progeny centriole may form adjacent to each parental centriole in the S phase (copy number control). In contrast to encouraging recent progress in understanding copy number control of centriole duplication in the S phase, the mechanisms licensing only one centriole duplication cycle within a cell cycle are not well understood. However, our study shows the great importance of CEP44 in controlling the centriole number during cell cycle progression, providing important insight into the detailed mechanisms underlying centriole duplication.

It also has been reported that CEP44 functions in centrosome cohesion by stabilizing rootletin (19). Centrosome cohesion is mediated by a loose connection between the proximal ends of the two parental centrioles, which is known as the G1-G2 tether (GGT). GGT assembly occurs in the G1 phase, which occurs simultaneously with, or just after, dissolution of the centriole engagement component termed the S-M linker (SML). Therefore, it has been speculated that GGT assembly depends on SML dissociation (4), but few studies have assessed the relationship between the SML and GGT. Apart from the function of CEP44 in centrosome cohesion, we found that CEP44 is involved in centriole engagement, suggesting the existence of some relationship between the SML and GGT, and additional studies will likely further illuminate the close relationship between the SML and GGT.

The localization and function of CEP44 led us to speculate that CEP44 may be a part of the SML. Previous studies found that SML functionality depends on the SAS-6–ANA2 complex (43) and suggested that cohesin might constitute part of the SML (44). However, the composition of the SML remains poorly understood. In this study, we found that CEP44 does not interact with the SAS6-STIL complex or cohesin (data not shown). Based on our results, it is plausible that the luminal protein CEP44 in the proximal part of the procentriole interacts with PCM protein CEP57/CEP57L1 in pre-existing centrioles to maintain centriole engagement, indicating that CEP44 is a potential SML component. Future studies aimed at determining whether CEP44 is a component of the SML will shed new light on the molecular mechanisms regulating centriole engagement.

We found that CEP44 is highly expressed in G2/M phase HeLa cells and nearly absent in MDA-MB-436 breast cancer cells, indicating that the protein level of CEP44 is precisely modulated during different stages of the cell cycle and in specific cell types. However, Aurora A-mediated phosphorylation of CEP44 had no effect on the protein level of CEP44 in our assay (Figure 6–figure supplement 1E). It is possible that epigenetic mechanisms or post-translational modifications function to regulate the protein level of CEP44.

Centrosome proteins have been reported to represent potent tumor suppressors or activators, because centrosome amplification is extremely common in a wide range of malignancies, including breast, prostate, ovarian, and lung tumors, and it is closely linked to tumor growth and a worse prognosis (45). Preclinical research has shown that extra centrosomes can directly cause tumor initiation, at least partly by counteracting centrosome functionality-decreasing mutations or by increasing chromosome instability through transiently formed abnormal spindles that result in missegregation because of deficiency in mitotic kinetochore-spindle attachments (46, 47). Therefore, due to insufficient CEP44, the presence of supernumerary centrosomes and abnormal spindle structure may compensate for impaired centrosome functionality of CCC during breast cancer initiation. Thus far, anti-mitotic drugs such as Aurora A and Plk1 inhibitors have largely failed in clinical contexts due to their low efficacy and serious side effects (48, 49). Notably, defective mitosis targeted by therapeutic strategies is always induced by centrosome amplification; therefore, it should be noted whether patients show centrosome amplification, and whether these inhibitors affect centrosome amplification, to ensure the achievement of a precise anti-tumor effect.

## Materials and Methods

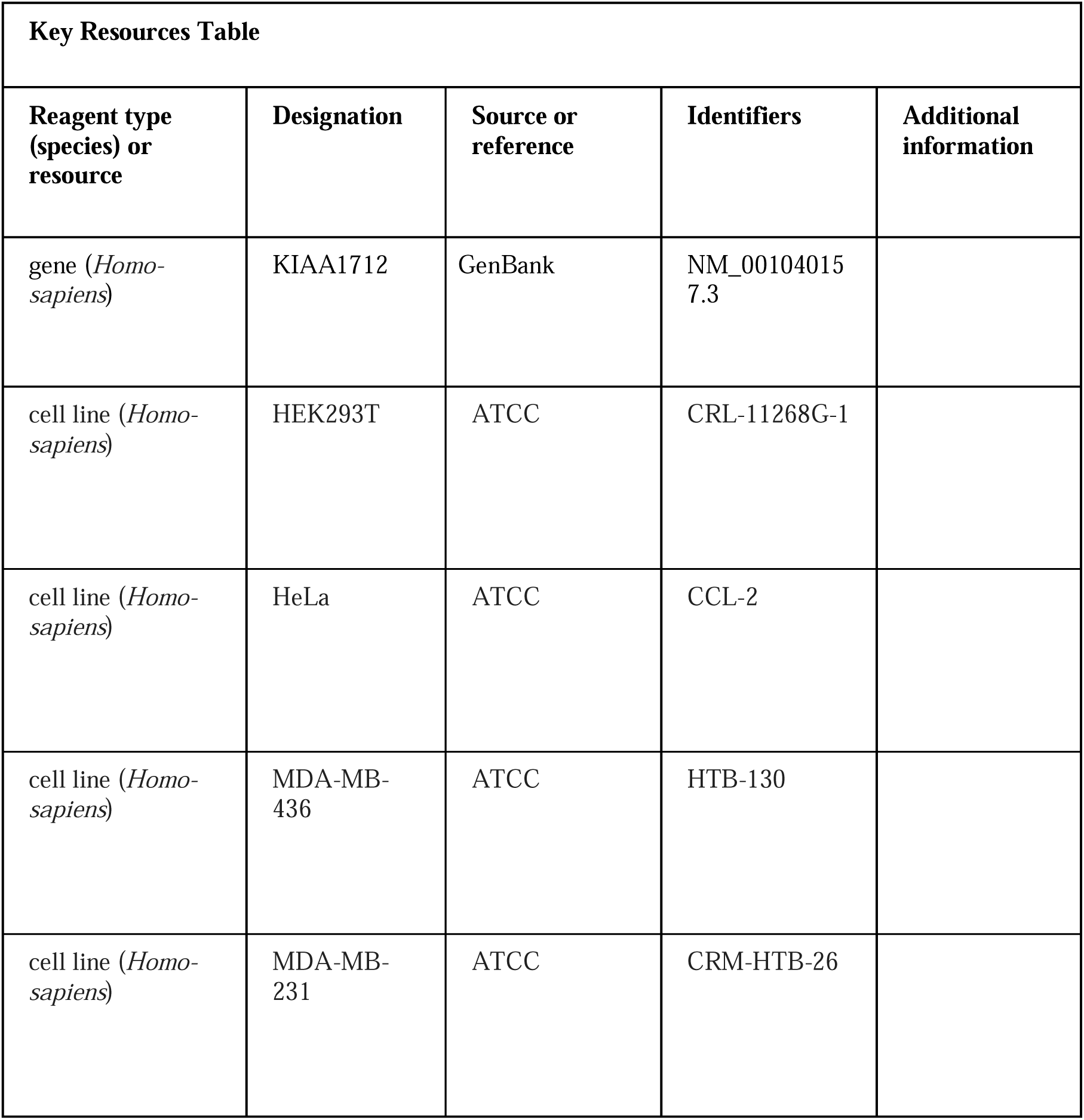
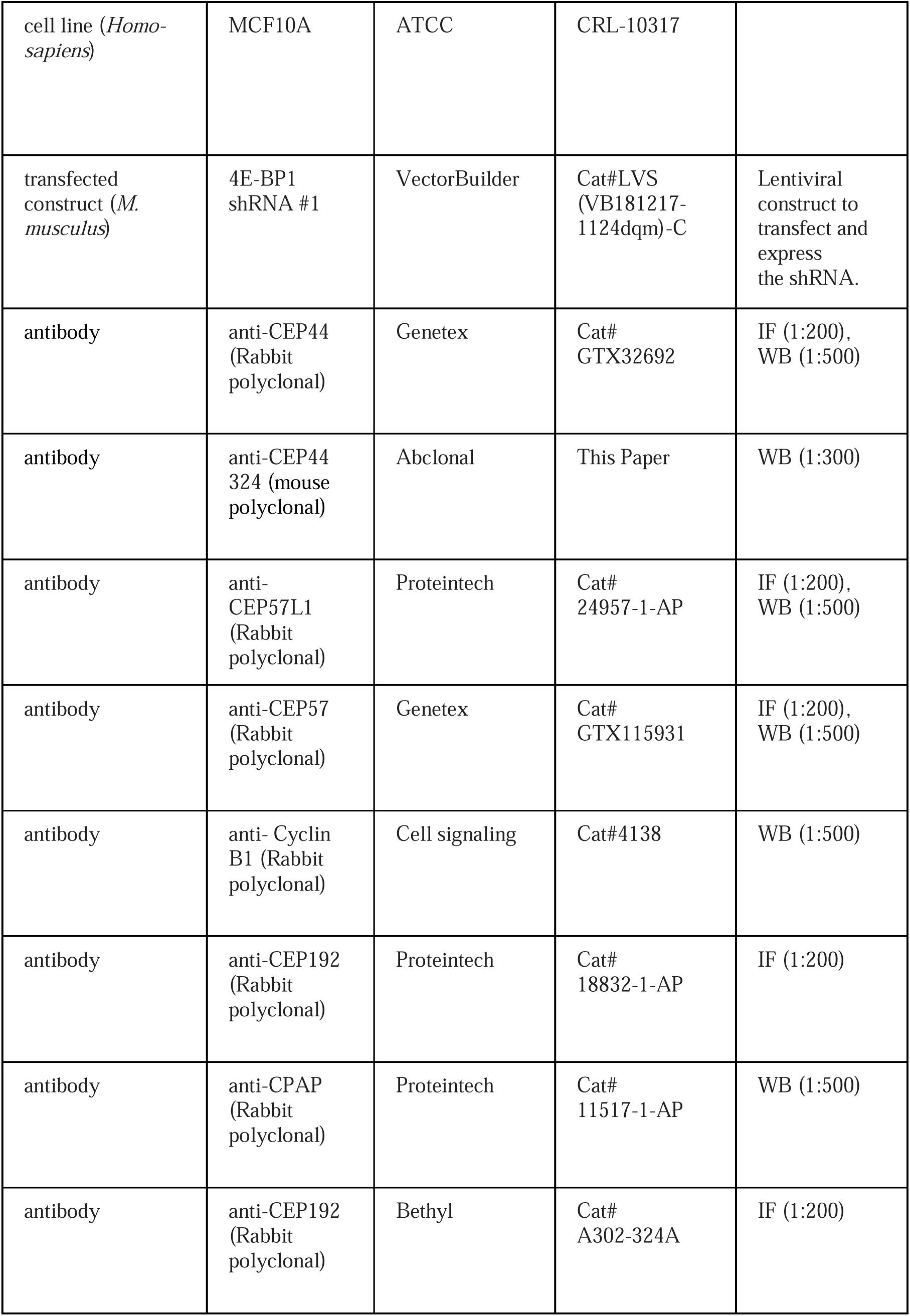
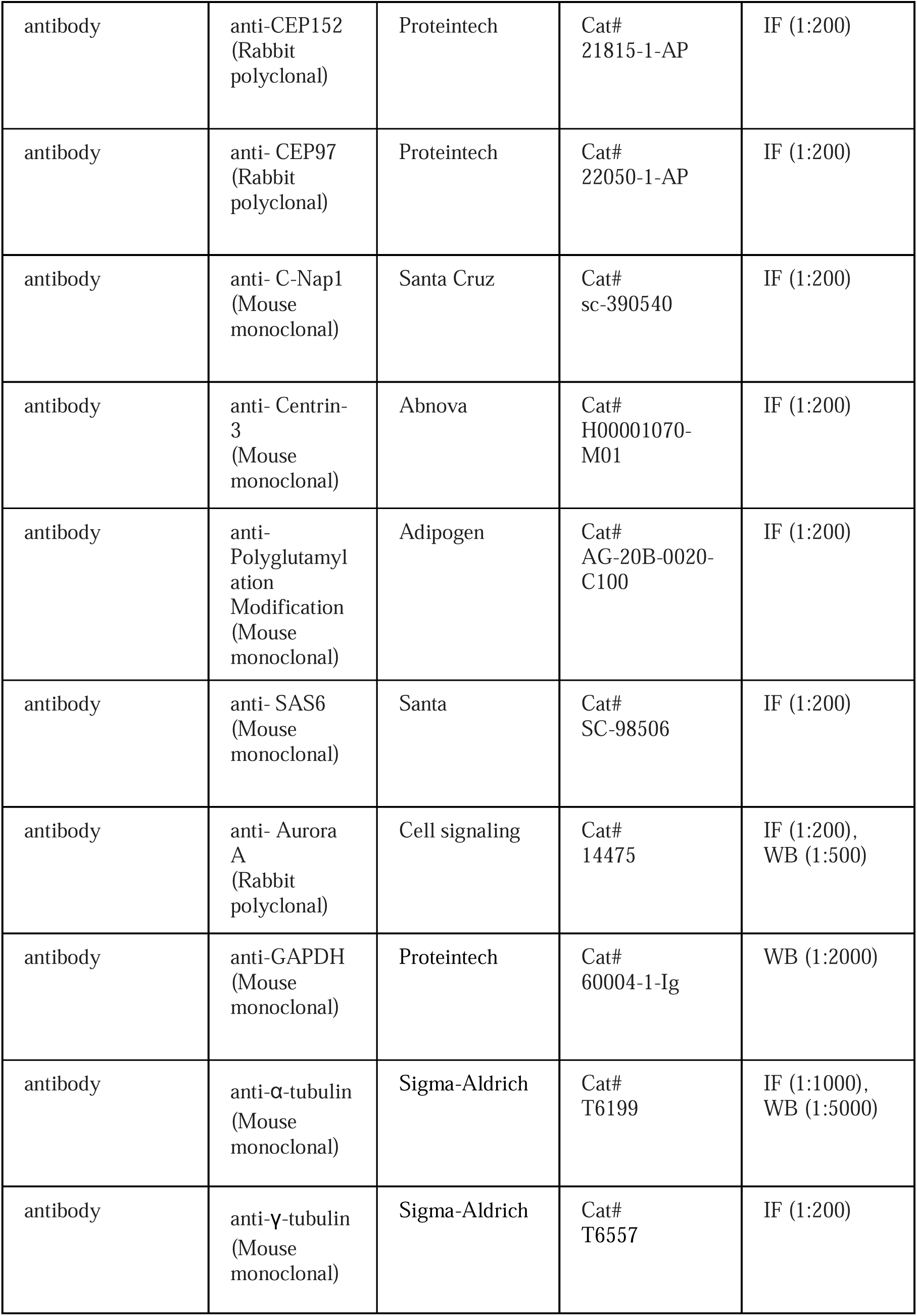
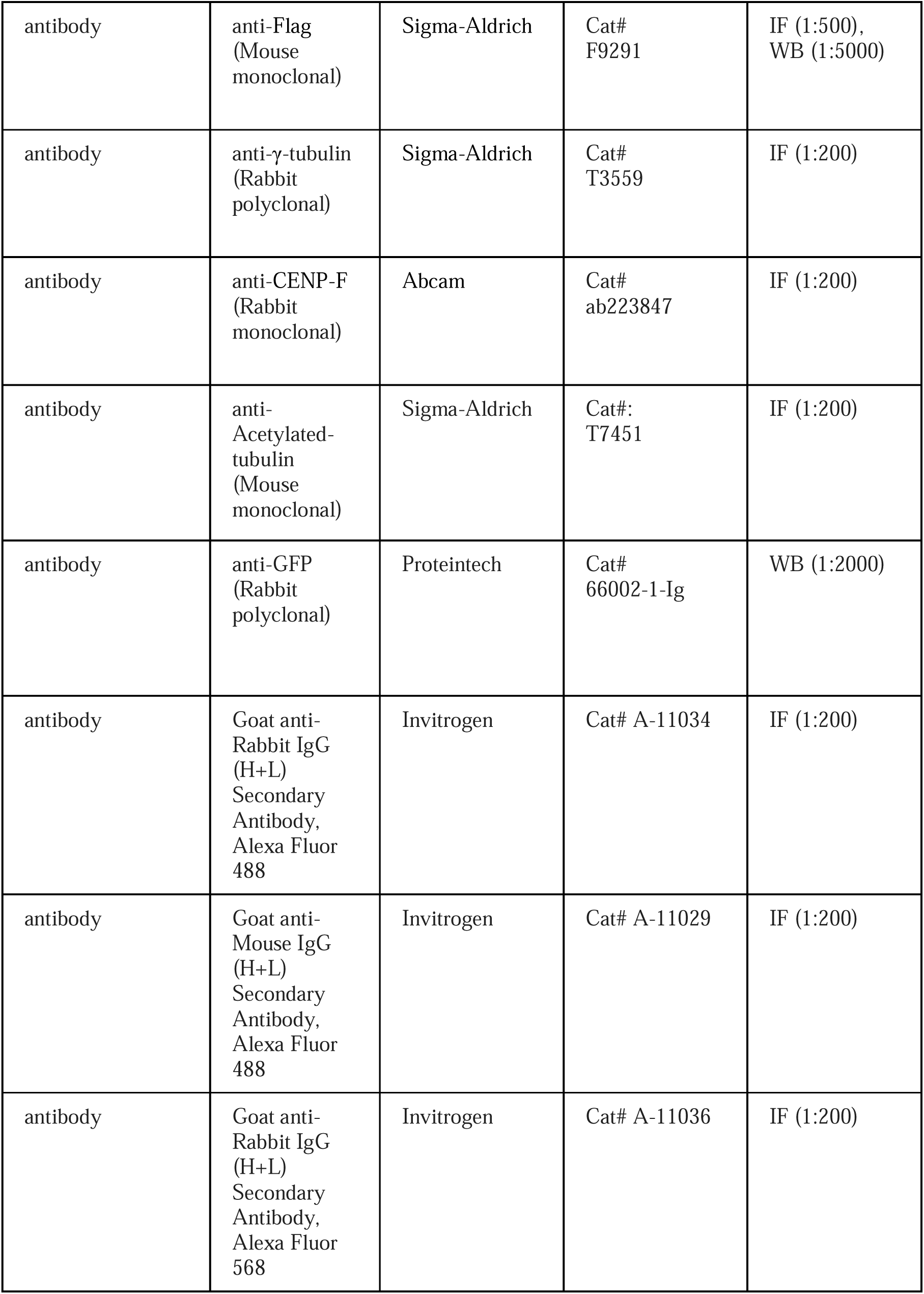
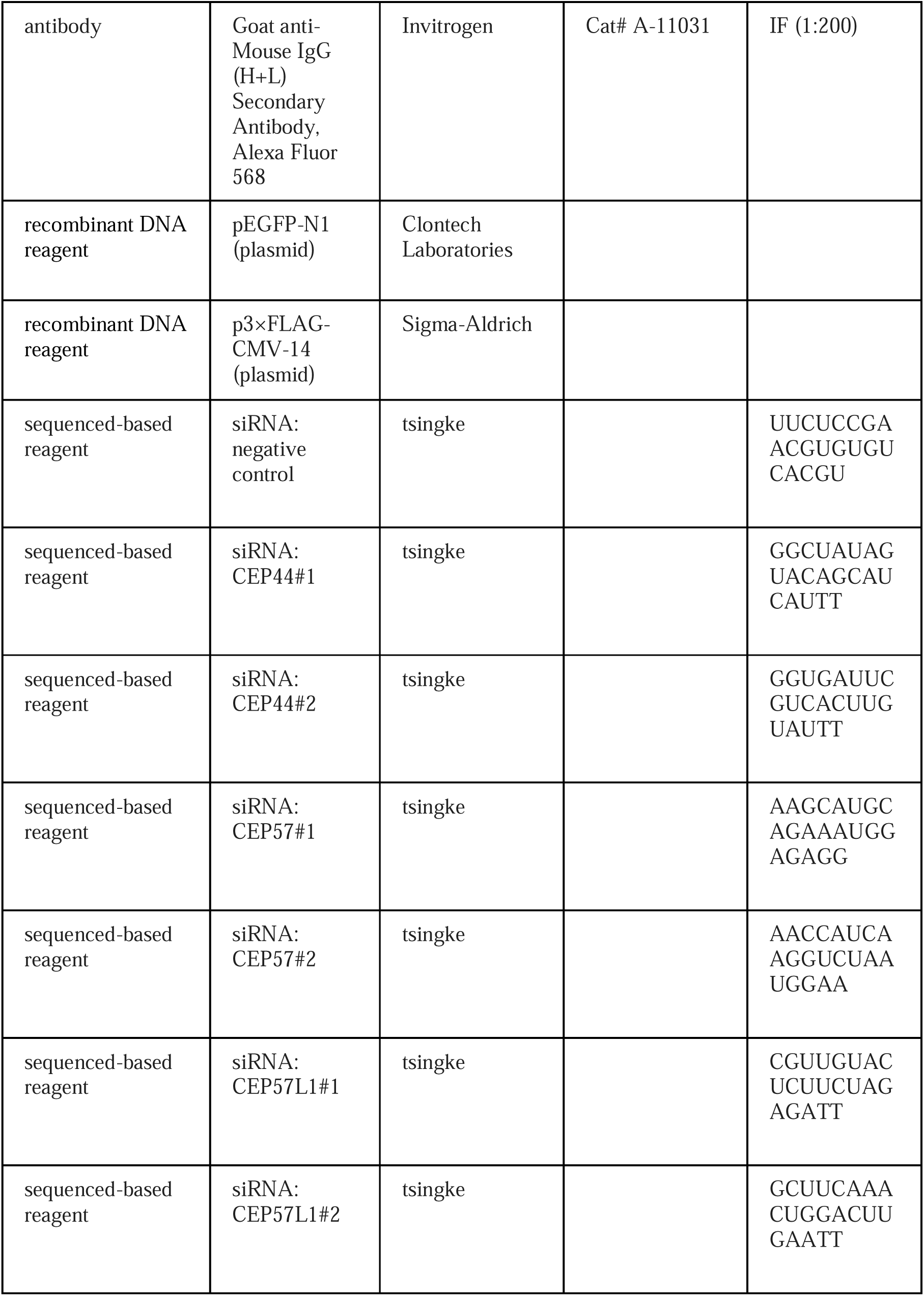
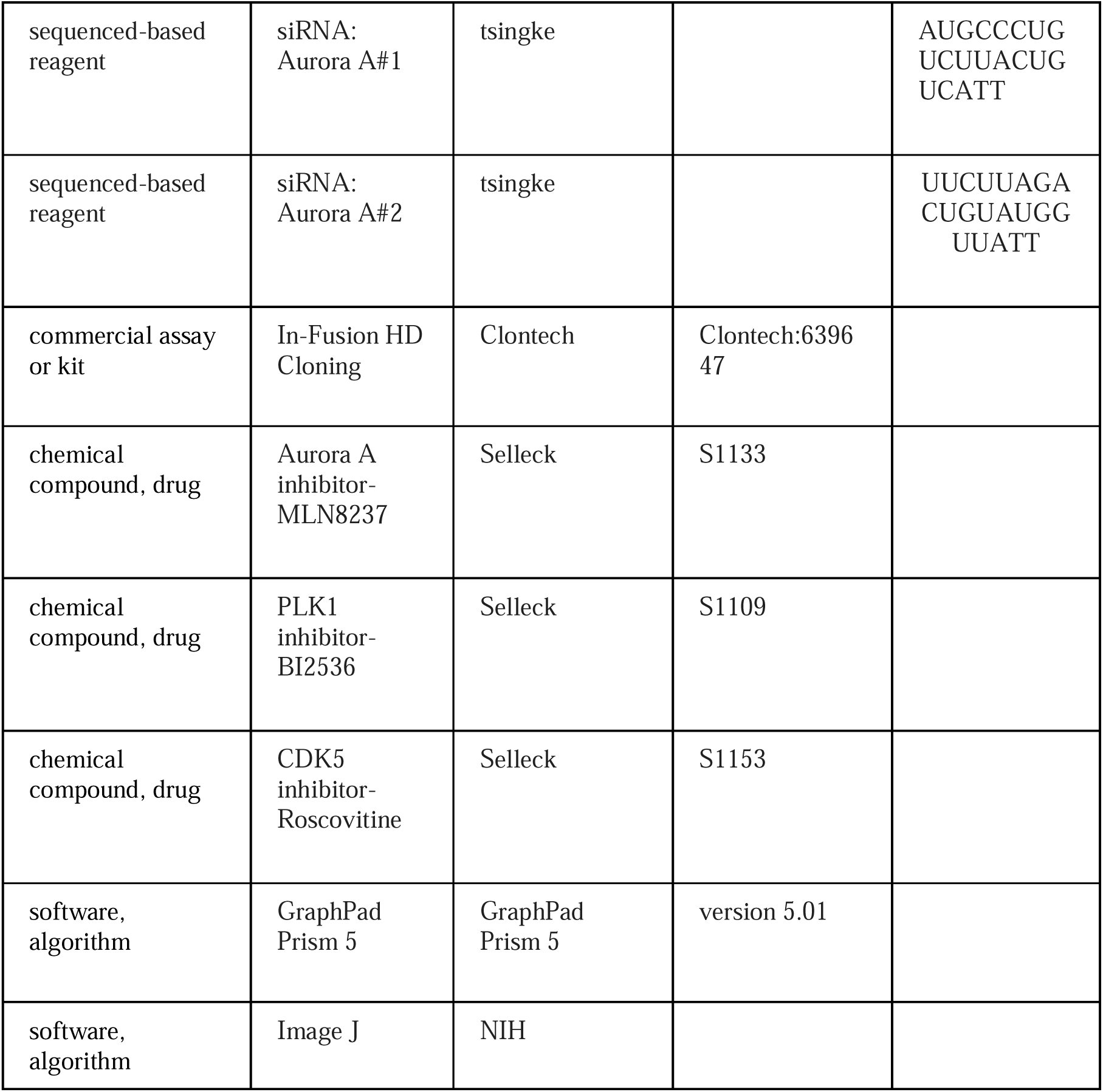

### Ethics statement

Breast tissue samples were collected during surgical treatment at the Oncology Department of the First Affiliated Hospital of Xinxiang Medical University (Henan, China). All samples were derived from idiopathic breast cancer patients, and none of these patients had received any treatment before surgery. These fresh resected biopsies of breast cancer were 4% formalin-fixed for immunohistochemical and pathological grading studies. Written informed consent for the samples obtained under protocols was approved by the Ethics Committee of the First Affiliated Hospital of Xinxiang Medical University (XYLL-2021B001). Diagnosis of all samples was confirmed by at least two independent pathologists.

### Plasmid construction

Human CEP44 (NM_001040157.3 [https://www.ncbi.nlm.nih.gov/nuccore/NM_001040157.3]), CEP44 fragments, and CEP44 mutants were amplified via PCR and cloned into pEGFP-N1 (Clontech Laboratories) and p3×FLAG-CMV-14 (Sigma-Aldrich). Other plasmids used in this study, including Centrin-3, Aurora A, Plk1, Nek2A, and CDK1 were generated as described previously (50).

### Antibodies

All antibodies used in this study are listed in the Key Resources Table. The mouse polyclonal antibody against CEP44 was generated using full-length human CEP44 fused with glutathione-S-transferase (GST). Anti-phosphorylated S324-CEP44 antibodies were raised in rabbits using a keyhole limpet hemocyanin (KLH)-conjugated DLLNPHRK(pS)EVERPAS peptide, which was affinity-purified on nonphosphorylated- and/or phosphorylated-epitope-bound columns (ABclonal Biotechnology).

### Cell culture and transfection

HEK293T (CRL-11268G-1), MCF10A, MDA-MB-436, MDA-MB-231and HeLa (CCL-2) cells were obtained from the American Type Culture Collection (ATCC). HEK293T (CRL-11268G-1), HeLa (CCL-2), MDA-MB-436 and MDA-MB-231 cells were cultured in DMEM (GIBCO) supplemented with 10% FBS (GIBCO or CellMax) at 37 °C under 5% CO2, with MDA-MB-436 cells adding 10 µg/ml insulin and 16 µg/ml glutathione. MCF10A was cultured in a base medium from Lonza/Clonetics Corporation (MEGM, Kit Catalog No. CC-3150) supplemented with 5% horse serum and 100 ng/ml cholera toxin. To avoid mycoplasma contamination, mycoplasma detection was implemented in these cell lines using GMyc-PCR Mycoplasma Test Kit (YEASEN, 40601ES20). Plasmids were transfected into HEK293T and HeLa cells using jetPEI (Polyplus transfection) or LipofectamineTM 3000 (Invitrogen), and siRNAs were transfected into HeLa cells using LipofectamineTM 2000, according to the manufacturer’s protocols.

### Cell synchronization and drug treatment

Synchronized HeLa cells in mitosis were treated with 100 ng/ml Nocodazole for 16-18 h. For double-thymidine blocking, HeLa cells were firstly subjected to 2.5 mM thymidine for 18-24 h, then released for 12 h, and subsequently blocked again for 18-24 h. To identify the kinase of CEP44, HeLa cells were mock-treated or treated with a kinase inhibitor: 100 nM MLN8237 (an Aurora A inhibitor) for 24 h, 100 nM BI2536 (a PLK1 inhibitor) for 4 h or 25 μM Roscovitine (a CDK5 inhibitor) for 18 h.

### Gene silencing by siRNA

All siRNAs used in this study were purchased from GenePharma. CEP57- and CEP57L1-siRNAs were designed according to a previous study (16, 51). The siRNA sequences are shown in Key Resources Table. The siRNA transfection procedure was performed for 48-72 h based on the manufacturer’s protocols. For the rescue assay, six silent mutations were introduced into the CEP44 sequence to construct siRNA-resistant CEP44 (resCEP44) (5’-GCTATAGTACAGCATCAT-3’ mutated to 5’-GATACAGCACCGCTTCATC-3’). HeLa cells were transfected with resCEP44 for 48 h after 24 h siRNA transfection.

### Western blotting

The samples were boiled at 100°C for 5 min in SDS loading buffer and then loaded in SDS-PAGE gels. The resolved samples were transferred to PVDF membranes (Millipore). The membranes were incubated with the primary antibodies listed in Key Resources Table, followed by incubation with HRP-conjugated secondary antibodies.

### λ-PPase treatment and Phos-tag assay

Cells cultured in a 35 mm dish were rinsed twice with tris-buffered saline (TBS) (50 mM Tris-HCl, 150 mM NaCl, pH 7.5) for collection, followed by lysis with RIPA buffer (1% NP-40, 150 mM NaCl, 0.5 mM EDTA, 0.5% sodium deoxycholate, 0.1% SDS, 50 mM Tris-HCl, 2 mM PMSF, protease inhibitor cocktail, pH 7.4) on ice for 30 min. After centrifugation at 12,000 × *g* for 15 min at 4 °C, the lysate supernatants were mixed with 10× NEBuffer Pack for Protein MetalloPhosphatases (PMP), 10× MnCl_2_, and 400 U λ-PPase to react at 30 °C for 30 min, and quenched by the addition of SDS loading buffer. To analyze phosphorylated protein samples, Phos-tag gel electrophoresis (10% separating gel) was applied based on the manufacturer’s protocol (Wako Pure Chemical Industries). The gel was supplemented with 50 μM MnCl_2_ and 25 μM Phos-tag Acrylamide AAL-107 and polymerized overnight. Samples were run at constant current of 30 mA per gel until the bromophenol blue arrived at the bottom of the separating gel. Before transfer of proteins, the gels were washed in 1 mM EDTA-containing transfer buffer for 30 min and then soaked in transfer buffer without EDTA for 20 min before the proteins were transferred to PVDF membranes for western blotting.

### Immunoprecipitation

HEK293T or HeLa cells were collected after washing twice in cold PBS and lysed in lysis buffer (50 mM HEPES, 250 mM NaCl, 5 mM EDTA, 0.1% NP-40, 1 mM DTT, 10% glycerol, 2 mM PMSF, protease inhibitor cocktail, pH 7.5) on ice for 30 min. Proteins were recovered by centrifugation at 12,000 × *g* for 15 min at 4 °C, and the supernatant was incubated with primary antibodies, protein G-Sepharose or protein A-Sepharose beads (GE Healthcare) for 2 h at 4 °C. The beads were washed 3-5 times with cold lysis buffer, boiled at 100 °C in SDS loading buffer for 5 min, and subjected to western blotting.

### Clinical samples and immunohistochemistry

Tissue microarray of primary HCC samples was obtained from Shanghai Outdo Biotech Co., Ltd. and US Biomax Inc. (Rockville, MD, USA). Immunohistochemical (IHC) staining was performed as the following steps40. Formalin-fixed, paraffin-embedded tissue slides were dewaxed with xylene and rehydrated by a graded series of alcohols, followed by antigen retrieval and block with 5% BSA for 60 min. Incubation was carried out at 4 °C for overnight with the primary antibody. Primary antibodies included: anti-TRIM25 polyclonal antibody (1:200; Abclonal), anti-KEAP1 polyclonal antibody (1:200; Proteintech), and anti-Nrf2 polyclonal antibody (1:200; Proteintech). Signals were detected using Envision-plus detection system (Dako, Carpinteria, CA, USA) and visualized following incubation with 3,3′-diaminobenzidine.

### Immunofluorescence and live cell imaging

Cells on coverslips were fixed using cold methanol for 10 min at −20°C and washed 3 times with cold PBS. Permeabilized cells were blocked with 4% Bovine Serum Albumin (BSA) and subsequently incubated with primary antibodies, followed by Alexa Fluor 488/568/647-conjugated secondary antibodies. Next, the cells were stained with 1 μg/ml DAPI. The samples were observed and photographed using a fluorescence microscope (TH4-200, Olympus) equipped with an Apo Oil Objective lens (60×, NA 1.42, Olympus), and images were taken using DP controller software (Olympus). Three-dimensional super resolution images were acquired using a three-dimensional structured illumination microscope (3D-SIM) equipped with an Apo Oil Objective lens (100×, NA 1.49, Nikon) (N-SIM System, Nikon), and images were taken and reconstructed to maximum projections using NIS-Elements AR software (Nikon). A confocal microscope (LSM-710 NLO, Zeiss) equipped with a 100× /1.4 NA objective lens and a super-resolution confocal microscope (Leica TCS SP8 STED × 3) equipped with a 100× /1.4 NA objective lens were used for cell observation. Images were taken using LAS X software (Leica), deconvolved using Huygens software (Scientific Volume Imaging, Netherlands), and processed using Photoshop (CC; Adobe). Cells transfected with H_2_B-RFP for 18 h were observed using a microscope (Nikon) with spinning-disk UltraView VoX system (PerkinElmer). Cells were cultured in a chamber with 5% CO_2_ at 37 L, and time-lapse images were taken and post-processed using Volocity software (PerkinElmer).

### Construction of CEP44 knockout cell lines

CEP44 knockout HeLa cell lines were constructed using the CRISPR/Cas9 approach (52). Target oligos were generated and ligated into U6-sgRNA vectors, and then transfected with pSpCas9 and pcDNA3.1-Puro vectors. Clonal isolation of puromycin-resistant HeLa cells was performed using flow cytometry analysis. To identify CEP44 knockout clones, cells were subjected to genome sequencing and western blotting analysis. The target oligos for gRNA are designed as 5’-AAAAGCAGTTTATCCAATGT-3’ and 5’-GGTAAGTCAGAACCTCCTTT-3’.

### Statistics

The RNA-sequencing (RNA-seq) data was obtained from TCGA. Kaplan-Meier analysis was performed for overall survival analyses and recurrence free survival. Overall survival was defined as time from the first treatment after primary diagnosis to death of any cause. Recurrence free survival was defined as time from the first treatment after primary diagnosis to death or recurrence of any cause. The target regions were drawn around the fluorescent dots and measured using ImageJ software (NIH). Either a two-tailed unpaired Student’s t-test or one-way ANOVA with Dunnett’s multiple comparisons test was applied using GraphPad Prism 5 (version 5.01; GraphPad, US) to compare differences between groups. The results are presented as the means ± S.E.M, and p < 0.05 was considered to indicate a statistically significant difference, unless otherwise stated.

## Data availability

All data needed to evaluate the conclusions in the paper are present in the paper.

## Acknowledgments

The authors thank C. Zhang (School of Life Sciences, Peking University) for providing the H2B-RFP plasmid; C. Shan (the Core Facilities of Life Sciences, Peking University) and J. Zhou (Nikon) for 3D-SIM and confocal microscopy imaging; L. Liu (Mass Spectrometry Facility of National Center for Protein Sciences at Peking University) for assistance with protein modification analysis; C. Yue (Department of Laboratory Medicine, Central People’s Hospital of Zhanjiang) for providing breast cancer cell lines; and Y. Qi (Department of Pathology, Central People’s Hospital of Zhanjiang) for observation of pathological tissue slices. The authors thank all members of the laboratory for helpful discussion. This work was supported by the National Natural Science Foundation of China (31830110, 32130024 and 32100542) and Zhanjiang Science and Technology Project (2022A01104).

## Author contributions

DHZ, JLT, NH and JGC designed the experiments. DHZ, WLW, FYL and MJY performed experiments. DHZ, WLW, FYL and MJY performed data analysis. FYL and MJY collected breast cancer tissue samples. DHZ, WLW and NH wrote and edited the manuscript. DHZ and JGC provided funding for this study. The order of the co-first authors was assigned on the basis of their relative contributions to the study.

---

**Figure 1–figure supplement 1.**
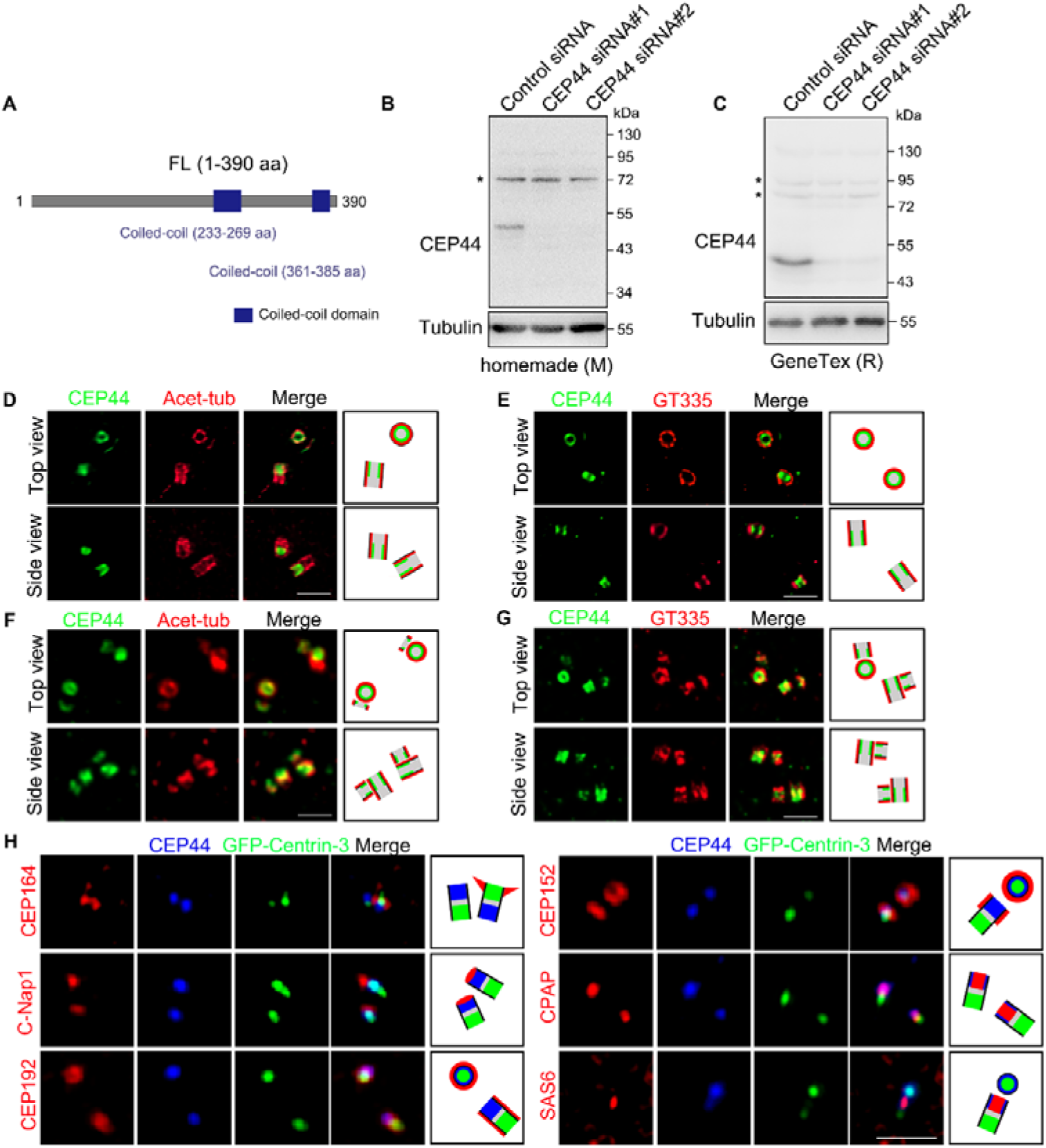
CEP44 is a centriole luminal protein localized at the proximal end. (A) Schematic of human CEP44. FL, Full length; Coiled-coil domains, blue. (B-C) Immunoblots of CEP44 in control and CEP44-depleted HeLa cells with a homemade (B) or commercial (C) antibody against CEP44. Tubulin was used as a loading control. Asterisk: non-specific band. (D-G) Immunostaining of CEP44 (green) with Acet-tubulin (red) (D and F) or GT335 (red) (E and G) in HeLa cells. Centrioles were duplicated in (F) and (G). (H) Immunostaining of CEP44 (blue) with CEP164, C-Nap1, CEP192, CEP152, CPAP or SAS6 (red) in CEP44-overexpressed HeLa cells. Bars: 500 nm (D-G), 2 μm (H).

**Figure 3–figure supplement 1.**
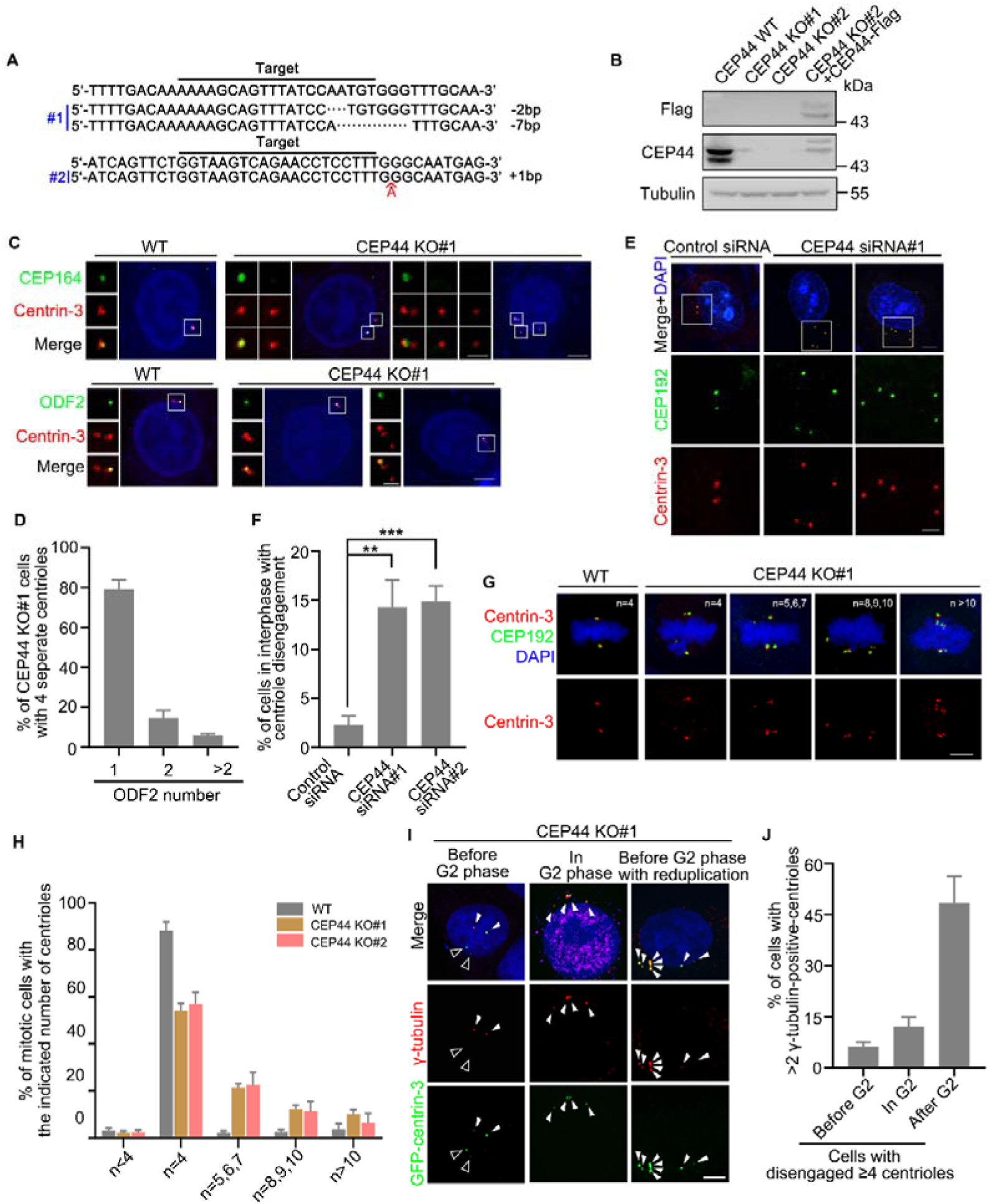
Depletion of CEP44 results in centriole disengagement and centriole overduplication. (A) Schematic of the edited sequence of CEP44 in CEP44 knockout HeLa cells. (B) Immunoblots of the indicated proteins in wild-type (WT), CEP44-knockout (KO) and CEP44-rescued HeLa cells. Tubulin was used as a loading control. (C) Immunostaining of Centrin-3 (red) with CEP164 (green) (upper) or ODF2 (green) (lower) in wild-type (WT) and CEP44-knockout (KO) HeLa cells. (D) Quantification of the indicated ODF2 number with the percentage of 4 separate centrioles in CEP44 KO HeLa cells. (E) Immunostaining of Centrin-3 (red) and CEP192 (green) in control and CEP44 knockdown HeLa cells. DNA was stained with DAPI (blue). (F) Quantification of the percentage of centriole disengagement in control and CEP44 knockdown HeLa cells in interphase. (G) Immunostaining of Centrin-3 (red) and CEP192 (green) in WT and CEP44 KO HeLa cells. DNA was stained with DAPI (blue). (H) Quantification of the percentage of mitotic cells with the indicated number of centrioles in WT and CEP44 KO cells. (I) Immunostaining of γ-tubulin (red) in CEP44 KO HeLa cells transfected with GFP-Centrin-3 (green). DNA was stained with DAPI (blue). The white and black arrowheads indicate the centriole dots co-labelled with or without γ-tubulin, respectively. (J) Quantification of the percentage of cells with more than 2 γ-tubulin-positive-centrioles in cells with disengaged ≥4 centrioles in the indicated cell cycle phase. For D, F and H, error bars represent the means ± S.E.M obtained for three independent experiments. \*\**p* < 0.01, \*\*\**p* < 0.001, as determined using one-way ANOVA with Dunnett’s multiple comparisons test. Bars: 5 μm (C, E, G and I).

**Figure 4–figure supplement 1.**
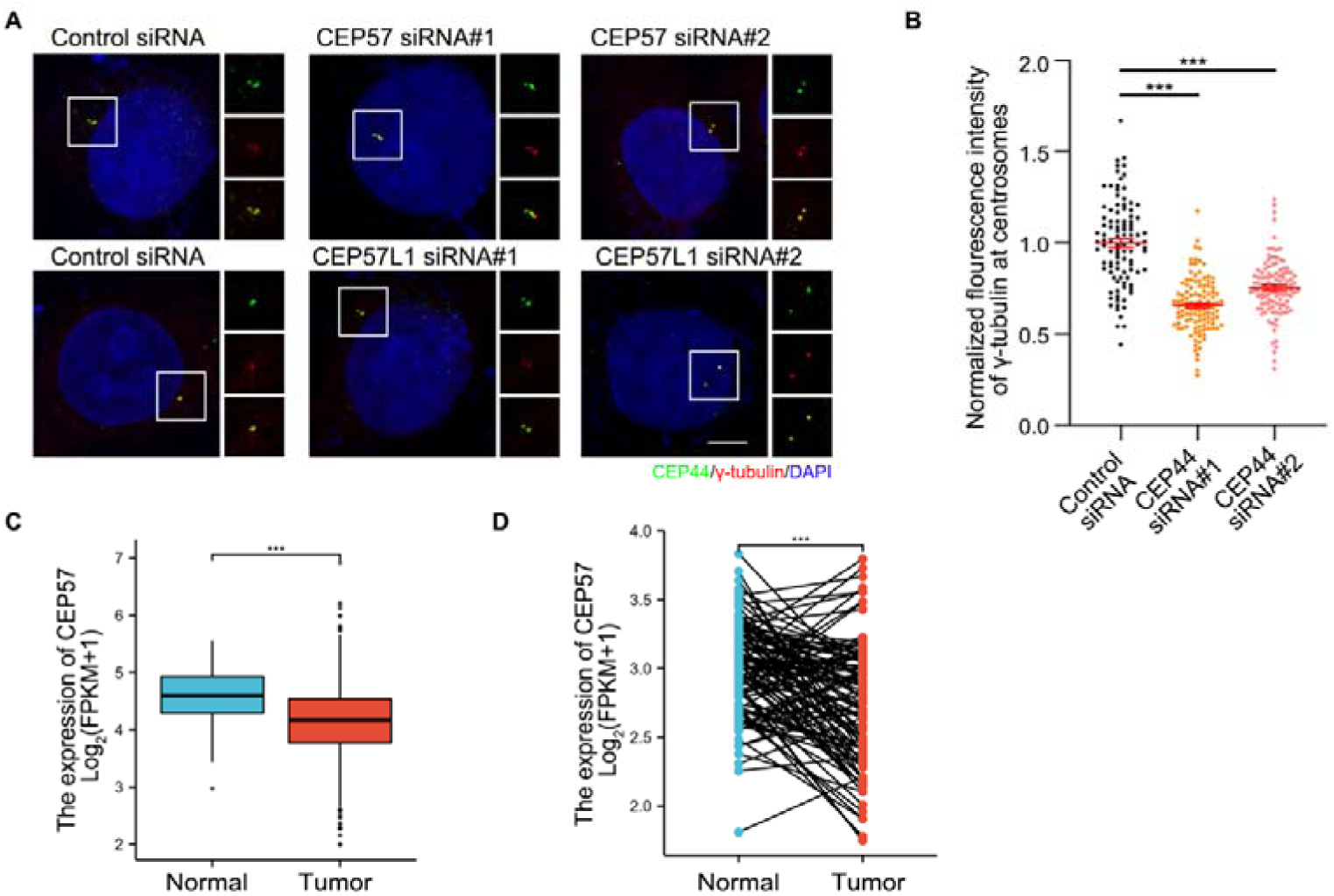
CEP57/CEP57L1 do not affect CEP44 localization at the centrosome. (A) Immunostaining of CEP44 (green) and γ-tubulin (red) in control and CEP44 knockdown HeLa cells. DNA was stained with DAPI (blue). (B) Quantification of the normalized fluorescence intensity of γ-tubulin at centrosomes in the cells from (A). (C) CEP57 transcript levels in normal human mammary tissues and breast carcinoma samples based on data sets from TCGA. (D) CEP57 transcript levels in paired tumor and tumor-adjacent tissue samples based on data sets from TCGA. For B, C and D, error bars represent the means ± S.E.M obtained for three independent experiments. \*\*\**p* < 0.001, as determined using one-way ANOVA with Dunnett’s multiple comparisons test. Bar: 5 μm (A).

**Figure 5–figure supplement 1.**
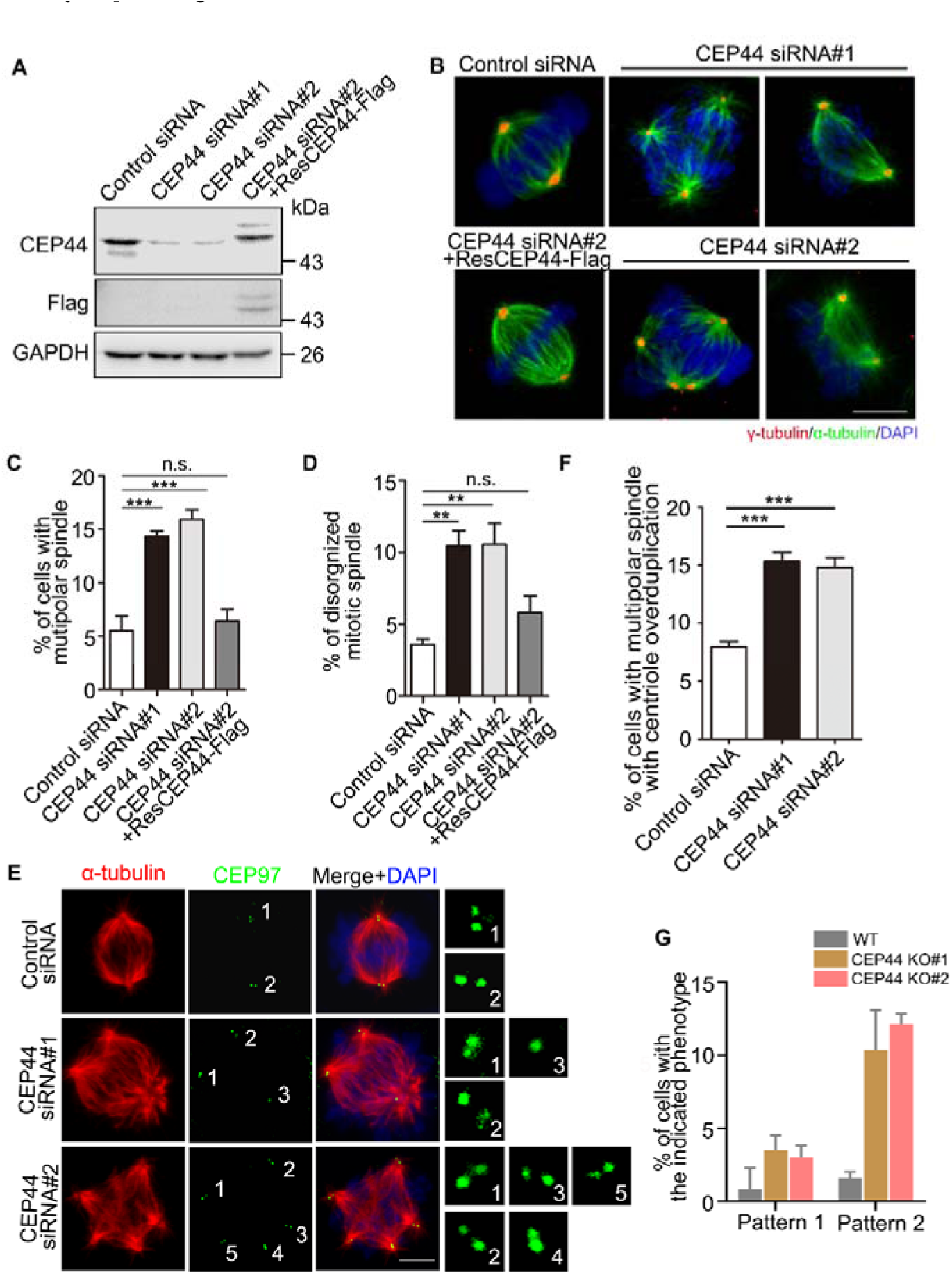
Knockdown of CEP44-centriole overduplication results in multipolar spindle. (A) Immunoblots of the indicated proteins in control, CEP44-depleted and CEP44-rescued HeLa cells. GAPDH was used as a loading control. (B) Immunostaining of α-tubulin (green) and γ-tubulin (red) in control, CEP44-depleted and CEP44-rescued HeLa cells. DNA was stained with DAPI (blue). (C) Quantification of the percentage of multipolar spindles in (B). (D) Quantification of the percentage of disorganized mitotic spindles in (B). (E) Immunostaining of α-tubulin (red) and CEP97 (red) in control and CEP44-depleted HeLa cells. DNA was stained with DAPI (blue). (F) Quantification of the percentage of multipolar spindles with centriole overduplication in (E). (G) Quantification of the percentage of pattern 1 and pattern 2 cells in WT and CEP44 KO cells. Pattern 1: premature centriole disengagement in interphase without centriole overduplication; Pattern 2: premature centriole disengagement in interphase with centriole overduplication. For C, D, F and G, error bars represent the means ± S.E.M obtained for three independent experiments. n.s., not significant, \*\**p* < 0.01, \*\*\**p* < 0.001, as determined using one-way ANOVA with Dunnett’s multiple comparisons test. Bars: 5 μm (B, E).

**Figure 6–figure supplement 1.**
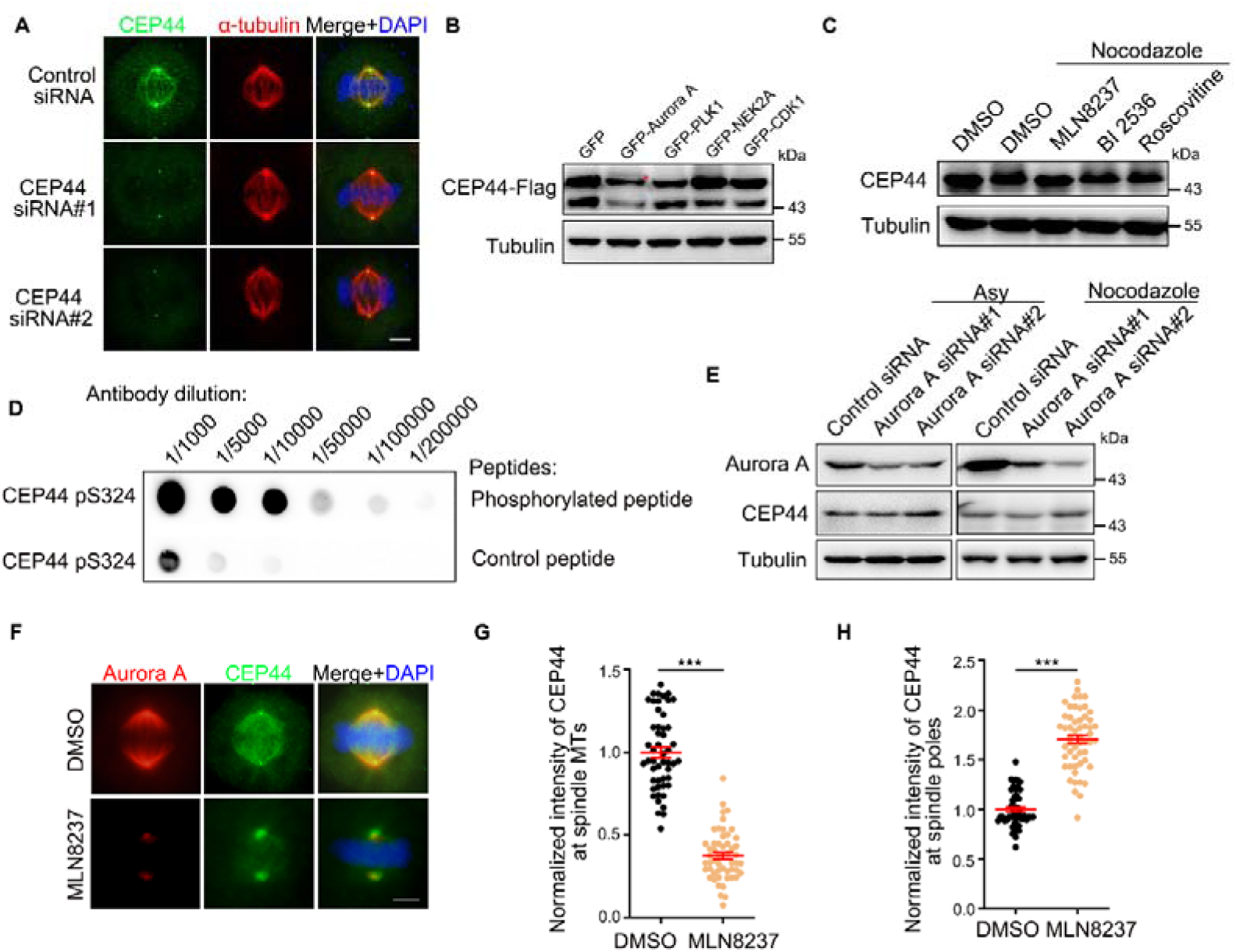
CEP44 phosphorylation affects localization and does not affect abundance. (A) Immunostaining of CEP44 (green) and α-tubulin (red) in control and CEP44 knockdown HeLa cells. DNA was stained with DAPI (blue). (B) Immunoblots of the lysates of CEP44-Flag-overexpressed HeLa cells with overexpression of the indicated kinases. Tubulin was used as a loading control. Asterisk: the shifted band. (C) Immunoblots of CEP44 in nocodazole-treated HeLa cells with the addition of the indicated inhibitors. Tubulin was used as a loading control. (D) Peptide competition assays using a phospho-CEP44 antibody against serine 324. (E) Immunoblots of Aurora A and CEP44 in control and Aurora A knockdown HeLa cells with or without nocodazole treatment. Tubulin was used as a loading control. (F) Immunostaining of Aurora A (red) and CEP44 (green) in control and MLN8237-treated HeLa cells. DNA was stained with DAPI (blue). (G) Quantification of the normalized intensity of CEP44 at spindle microtubules from (F). (H) Quantification of the normalized intensity of CEP44 at spindle poles from (F). For G and H, error bars represent the means ± S.E.M obtained for three independent experiments. \*\*\**p* < 0.001, as determined using Student’s t-test. Bars: 5 μm (A and F).

**Figure 7–figure supplement 1.**
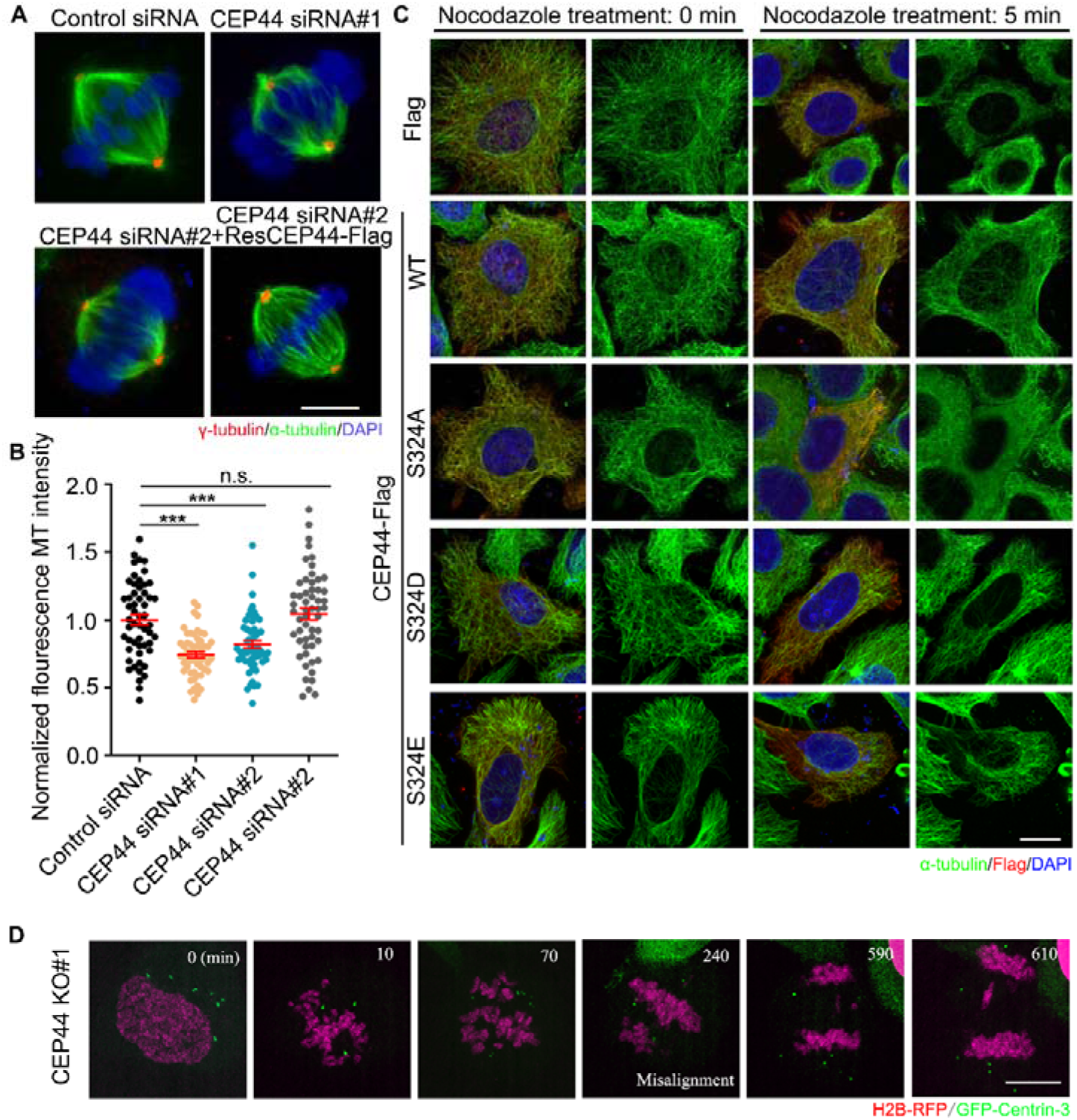
CEP44 is critical for spindle structure formation. (A) Immunostaining of γ-tubulin (red) and α-tubulin (green) in control, CEP44-depleted and CEP44-rescued HeLa cells. DNA was stained with DAPI (blue). (B) Quantification of the normalized fluorescence intensity of MT in the groups from (A). (C) Immunostaining of α-tubulin (green) and Flag (red) in HeLa cells expressing the indicated proteins with or without nocodazole treatment. DNA was stained with DAPI (blue). (D) Time-lapse images of cells stably expressing H2B-RFP and GFP-Centrin-3 in mitotic CEP44 knockout HeLa cells. For B, error bars represent the means ± S.E.M obtained for three independent experiments. n.s., not significant, \*\*\**p* < 0.001, as determined using one-way ANOVA with Dunnett’s multiple comparisons test. Bars: 5 μm (A, C and D).

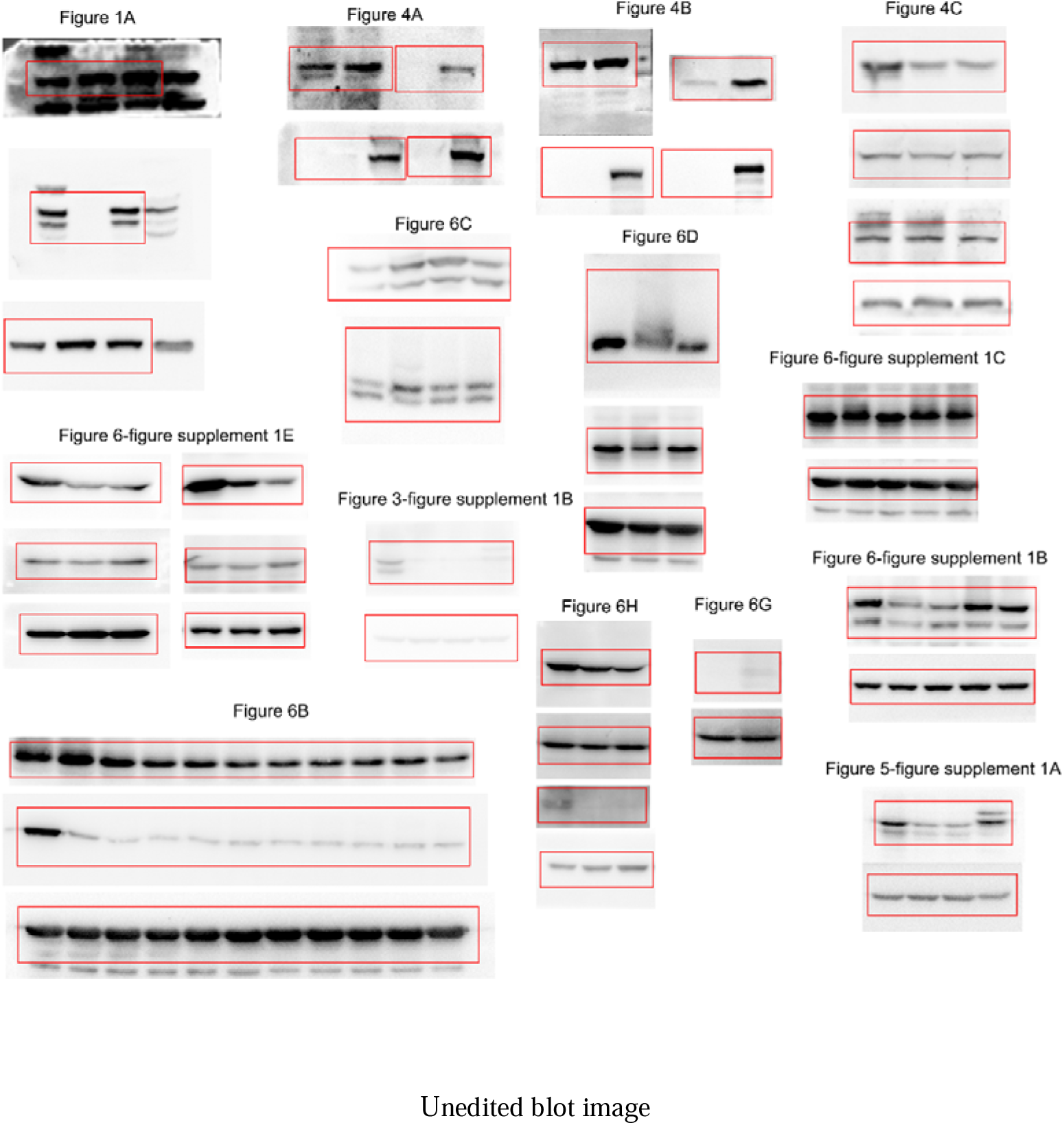

## Notes

**Conflict of interest:** The authors have declared that no conflict of interest exists.

https://elifesciences.org/reviewed-preprints/94405#tab-content

### Competing Interest Statement

The authors have declared no competing interest.

### Summary of Updates

Figure 3 figure supplement 1,Figure 5, Figure 5 figure supplement 1, Figure 7 figure supplement 1 and Figure 8 revised; Supplemental files updated.

