## Supplementary material for "CEP44 is required for maintaining centriole duplication and spindle integrity": Figure 1-source data 1

**Figure 1-source data 1.** Breast cancer patients information

|  | **Age** | **Gender** | **Clinical Stage** |
| --- | --- | --- | --- |
| 1 | 78.00 | female | T2N1M0 |
| 2 | 70.00 | female | T3N3M0 |
| 3 | 61.00 | female | T2N0M0 |
| 4 | 50.00 | female | T2N3M0 |
| 5 | 37.00 | female | T1N1M0 |
| 6 | 60.00 | female | T2N2M0 |
| 7 | 36.00 | female | T1N1M0 |
| 8 | 42.00 | female | T1N0M0 |
| 9 | 62.00 | female | T2N0M0 |
| 10 | 41.00 | female | T2N0M0 |
| 11 | 55.00 | female | T1N2M0 |
| 12 | 34.00 | female | T2N3M0 |
| 13 | 53.00 | female | T1N0M0 |
| 14 | 60.00 | female | T1N0M0 |
| 15 | 47.00 | female | T1N0M0 |
| 16 | 55.00 | female | T1N2M0 |
| 17 | 66.00 | female | T2N1M0 |
| 18 | 57.00 | female | T1N2M0 |
| 19 | 45.00 | female | T1N0M0 |
| 20 | 39.00 | female | T2N1M0 |
| 21 | 45.00 | female | T1N0M0 |
| 22 | 51.00 | female | T2N0M0 |
| 23 | 49.00 | female | T2N0M0 |
| 24 | 48.00 | female | T2N0M0 |
| 25 | 53.00 | female | T2N0M0 |
| 26 | 45.00 | female | T1N0M0 |
| 27 | 58.00 | female | T2N0M0 |
| 28 | 56.00 | female | T3N0M0 |
| 29 | 49.00 | female | T2N0M0 |
| 30 | 51.00 | female | T2N1M0 |
| 31 | 53.00 | female | T2N2M0 |
